# Biphasic JNK–Erk Signaling Separates Induction and Maintenance of Cell Senescence after DNA Damage

**DOI:** 10.1101/2022.06.15.496288

**Authors:** Tatiana S. Netterfield, Gerard J. Ostheimer, Andrea R. Tentner, Peter K. Sorger, Kevin A. Janes, Douglas A. Lauffenburger, Michael B. Yaffe

## Abstract

Genotoxic stress in mammalian cells, including that caused by anti-cancer chemotherapy, can induce temporary cell cycle arrest, DNA damage-induced senescence (DDIS) or apoptotic cell death. Despite obvious clinical importance, it is unclear how the signals emerging from DNA damage are integrated together with other cellular signaling pathways monitoring the cell’s environment and/or internal state to control these different cell fates. Here, using a combination of single cell-based signaling measurements and tensor PLSR/PCA computational approaches, we show that the JNK and Erk MAPK signaling pathways regulate the initiation of senescence through the transcription factor AP-1 at early times after extrinsic DNA damage, and the Senescence Associated Secretory Phenotype, a hallmark of DDIS, at late times after damage. These results identify a time-based separation of function for the same signaling pathways beyond the classic DNA damage response that control the cell senescence decision and modulate the tumor microenvironment following genotoxic stress, and reveal a fundamental similarity between signaling mechanisms responsible for oncogene-induced senescence and senescence caused by extrinsic DNA damaging agents.

## INTRODUCTION

Eukaryotic cells recognize and respond to DNA damage by activating an evolutionarily conserved set of signaling pathways that are essential for maintaining genomic integrity and preventing cancer (Hoeijmakers, 2001; Jackson & Bartek, 2009). These DNA Damage Response (DDR) signaling pathways regulate DNA damage induced-cellular activities and outcomes including DNA repair, cell cycle arrest, senescence and apoptosis (Harper & Elledge, 2007; Matt & Hofmann, 2016). Cellular senescence and apoptosis are actively regulated cellular responses that reduce the likelihood of cancer by preventing cells with genomic damage (or cells at risk of genomic damage) from proliferating (Bartkova et al., 2006; D’Adda Di Fagagna, 2008; Gorgoulis et al., 2005; Van Nguyen, Puebla-Osorio, Pang, Dujka, & Zhu, 2007). Mutations and/or acquired defects that compromise the function of these DDR pathways result in enhanced mutagenesis, and underlie the development and progression of cancer (Ciccia & Elledge, 2010; Halazonetis, Gorgoulis, & Bartek, 2008; Kastan & Bartek, 2004).

In the canonical DDR signaling pathway, double stranded breaks (DSBs) in DNA stimulate the kinase activity of ATM that phosphorylates and recruits a suite of proteins, including the histone variant H2AX, thereby creating detectable foci of DNA damage response proteins in the area adjacent to the DSB (Furuta et al., 2003; Harper & Elledge, 2007; Hoeijmakers, 2001; Jackson & Bartek, 2009). ATM effectors include the checkpoint kinases Chk2, Chk1, and MK2, and the multi-functional transcription factor p53, which together communicate DNA damage to the cellular machinery responsible for cell cycle arrest and the induction of programmed cell death (Reinhardt & Yaffe, 2009). p53 is a central node in the DDR signaling network that contributes to transient cell cycle arrest and senescence by up-regulating the cyclin-dependent kinase inhibitor p21^Waf1^, and to apoptosis by transactivation of pro-apoptotic Bcl-2 protein family members (Childs, Baker, Kirkland, Campisi, & van Deursen, 2014; He et al., 2005; Levine & Oren, 2009; Toshiyuki & Reed, 1995). Tumor cells often have mutations in DDR components, including p53, which allows evasion of normal cell cycle control mechanisms and contributes to genomic instability. However, such defects can also sensitize tumor cells to killing and/or cell cycle arrest and senescence by classical DNA-damaging agents, such as ionizing radiation and chemotherapy drugs used to treat cancer (Helleday, 2008; Helleday, Petermann, Lundin, Hodgson, & Sharma, 2008; R. D. Kennedy & D’Andrea, 2006; Powell & Bindra, 2009).

Cellular senescence is a “catch-all” term that refers to three classes of irreversible cell cycle arrest – replicative senescence (RS), oncogene-induced senescence (OIS) and DNA damage induced senescence (DDIS) (Campisi & D’Adda Di Fagagna, 2007). RS occurs after eukaryotic cells have undergone sufficient rounds of replication to expose unprotected telomeric DNA, resembling an un-repairable DNA double strand break (DSB). Subsequent DDR signaling then results in permanent cell cycle arrest (D’Adda Di Fagagna et al., 2003; Herbig, Jobling, Chen, Chen, & Sedivy, 2004; Shay & Wright, 2004). OIS occurs when oncogene expression results in inappropriately high levels of proliferation, leading to DNA replication stress and collapsed replication forks. The resulting DSBs induce a DDR-dependent permanent cell cycle arrest (Bartkova et al., 2006; Halazonetis et al., 2008). DDIS occurs after exposure to sub-apoptotic, “intermediate” levels of DNA damage, including those caused by ionizing radiation (IR), platinum drugs, or topoisomerase inhibitors, resulting in damage that is too high for complete repair and cell cycle re-entry, but not high enough to induce cell death (Childs et al., 2014; D’Adda Di Fagagna, 2008). Senescent cells of all three classes are viable, metabolically active, enlarged and/or flattened in morphology, and strongly positive for cyclin dependent kinase inhibitors (CDKIs), persistent DNA damage induced foci (PDDF) and senescence-associated heterochromatic foci (SAHF) (Rodier & Campisi, 2011). In addition, senescent cells secrete a panel of inflammatory cytokines featuring high levels of IL-6 and IL-8 (Basisty et al., 2020; Coppé, Desprez, Krtolica, & Campisi, 2010; Coppé et al., 2008; Davalos, Coppe, Campisi, & Desprez, 2010; Rodier et al., 2009). Campisi and co-workers showed that this senescence-associated secretory phenotype (SASP) requires signaling through the DDR (Rodier et al., 2009) and p38/NF-кB pathways (Freund, Patil, & Campisi, 2011). Although DDR signaling appears required for driving cell senescence, the contribution of cytokine signaling pathways to cell fate decisions has not been as clearly defined.

The commitment of DNA damaged cells to transient cell cycle arrest, senescence, or apoptotic cell fates likely involve integrating DDR signaling with additional signaling pathways governing general stress and survival responses such as the Akt pathway, the NF-kB pathway, and the stress- and mitogen-activated protein kinase (SAPK/MAPK) pathways. Since cell stress and extracellular signals are both transduced through these pathways, they may serve as information processing junctions that integrate signals from the microenvironment with DDR signaling. Previous studies have indicated roles for the ERK, JNK and p38 SAPK/MAPK pathways in cell fate determination after DNA damage (Janes et al., 2006; M. J. Kim et al., 2008; Manke et al., 2005; Spallarossa et al., 2010; Tentner et al., 2012). However, the relative importance of these additional signal transduction pathways, and the manner and timing by which their signals are integrated together with those from the canonical DDR pathways to control the outcome of DNA-damaged cells, is not well understood. Therapeutic re-wiring of these pathways could allow control over cellular outcomes and improv the clinical response of tumors to canonical genotoxic therapies (Kong et al., 2020; Lee et al., 2012; Mokim Ahmed & Jian Li, 2007; Reinhardt, Aslanian, Lees, & Yaffe, 2007; Roberts & Der, 2007).

To investigate cell fate determination after DNA damage in a systematic manner, we undertook a quantitative time-resolved cell signaling and phenotypic response study in U2OS osteosarcoma cells exposed to different levels of doxorubicin-induced DNA damage with the intention of using data-driven models to nominate novel relationships between signaling activity and cellular outcomes. We were particularly interested in identifying signaling events that denoted different temporal stages on the paths to senescence and apoptosis. Using a structured, multidimensional dataset of signaling and DNA damaged-induced responses modeled by tensor-and matrix-factorization approaches, we now report that the transcription factor AP-1, and its upstream activators, the stress and mitogen activated protein kinases JNK and ERK, play important roles in the cell fate decision to senesce at arrest-inducing (but non-lethal) doses of DNA damage at early times, and contribute to the DNA damage induced SASP at later times. Taken together this systems-based analysis points to a fundamental role for SAPK/MAPK signaling in the regulation of DDIS and the SASP along separable distinct timescales, and reveals similarities between the signaling events responsible for both OIS and DDIS.

## RESULTS

### A Cue-Signal-Response Framework for Interrogating Cell Fate Choice After DNA Damage

DNA damaging agents induce cells to undergo cell cycle arrest, followed by either DNA repair and cell cycle re-entry, DNA damage induced senescence (DDIS) or apoptotic cell death. To quantitatively map the DNA damage cue-signal-response landscape (Janes et al., 2004), we systematically profiled human U2OS osteosarcoma cells treated with a range of doxorubicin doses (Fig. 1A). Doxorubicin—a commonly used chemotherapeutic agent used to treat a variety of human malignancies including osteosarcoma—induces DNA double-strand breaks (DSBs) primarily by inhibiting topoisomerase II, but is also known to generate reactive oxygen species (Gewirtz, 1999) and act as an intercalating agent (Bodley et al., 1989). U2OS cells were selected for study because they are widely used in studies of DNA damage signaling (Maki & Howley, 1997; Manke et al., 2005; Reinhardt et al., 2007), express wild-type p53, and undergo a full range of DNA damage induced cellular responses, including p53-dependent cell cycle arrest and apoptosis. U2OS cells do not express the cyclin-dependent kinase inhibitor p16, because their *INK4* locus has been silenced by methylation; nonetheless, this cell line is fully capable of undergoing senescence (Bartkova et al., 2006; Park et al., 2002).

**Figure 1.**
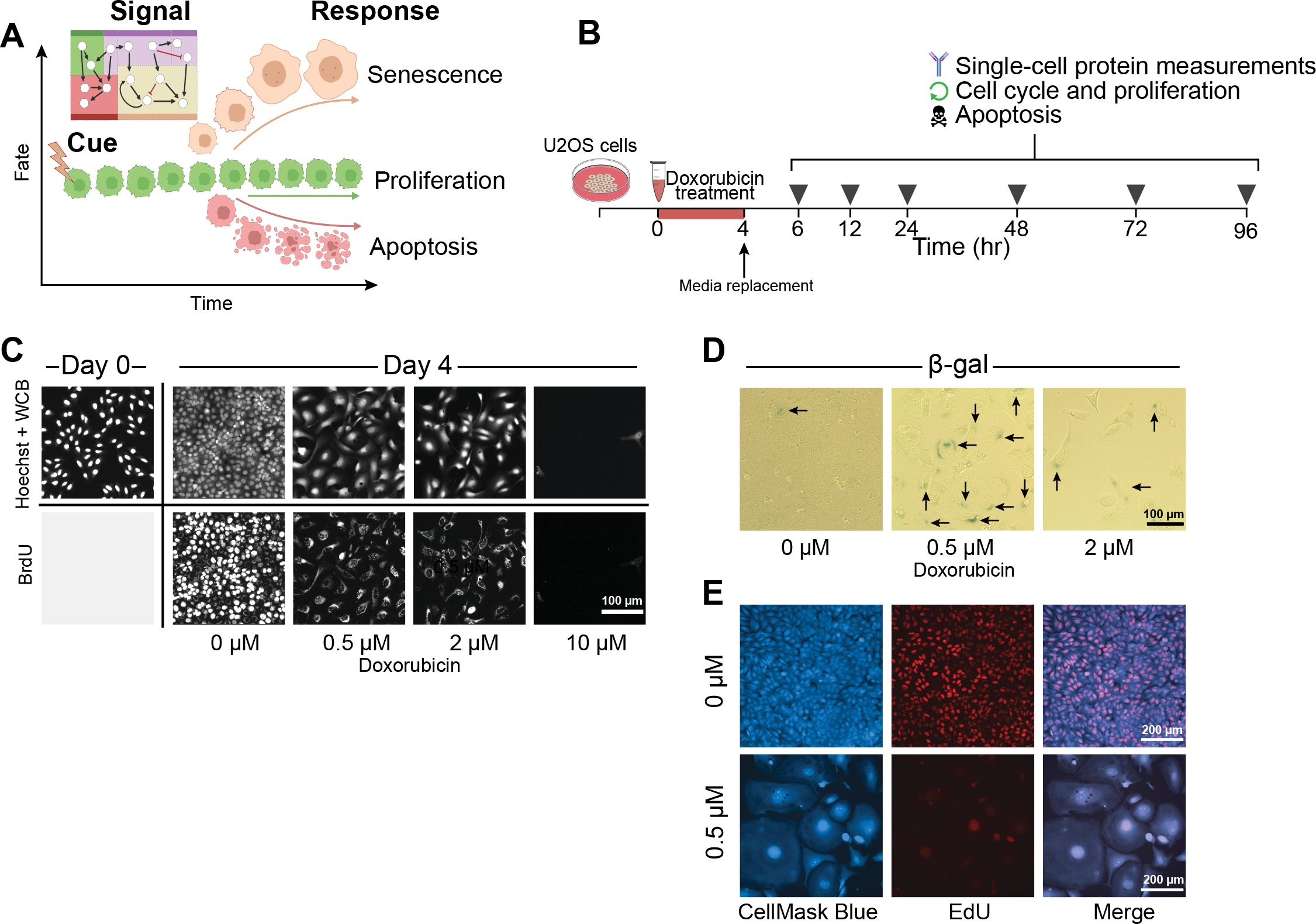
A cue-signal-response framework for cell fate decisions after DNA damage. **A)** Cartoon of cell fates of interest vs. time after DNA damage, created with BioRender.com. **B)** A schematic of how the signaling and response data was collected, using icons created in BioRender.com. For details, see text. **C)** U2OS cells were treated with a 4 hour pulse of either with DMSO (0 µM) or 0.5, 2, or 10 µM doxorubicin, and fixed 4 days after treatment. Hoechst and Whole Cell Blue (WCB) dye were used to visualize cell morphology, while immunofluorescence for BrdU DNA incorporation was used to label proliferating cells. **D)** β-galactosidase activity was measured by colorimetric staining in fixed U2OS cells 4 days after doxorubicin treatment. Black arrows denote positively stained cells. **E)** Cells were either treated with DMSO (0 µM) or with 0.5 µM doxorubicin, and then fixed 6 days later. Cell morphology was visualized with HCS CellMask Blue, and proliferation with EdU DNA incorporation. CellMask Blue channel was processed with a gamma of 0.5 to better visualize the cytoplasmic compartment.

U2OS cells were treated with 0.5, 2 or 10 µM doxorubicin or carrier for 4 hours, followed by media replacement. Individual cells were assayed for signaling events and phenotypic outcomes at 6, 12, 24, 48, 72 and 96 hours after the start of treatment (Fig. 1B). In addition, cell morphology was examined by Whole Cell Blue staining, and cell proliferation assessed by BrdU labeling and immunohistochemical detection at 96 hours after treatment. Mock treated U2OS cells actively proliferated during the 4 days after treatment as shown by increased cell density and BrdU incorporation (Fig. 1C). In contrast, both the 0.5 and 2 µM doxorubicin treatments arrested proliferation, as shown by the absence of nuclear BrdU incorporation. These doses of doxorubicin induced a change in morphology that is consistent with cellular senescence—cell size and nuclear size increased over the 4 day time course, and the cells assumed a ‘fried egg’ appearance (Amtmann, Eddé, Sauer, & Westphal, 1990). Interestingly, non-proliferating cells showed perinuclear BrdU staining after 24 hours of exposure to BrdU, likely resulting from BrdU incorporation into RNA (Wansink et al., 1993) that suggests metabolically activity is retained even though the cells are not proliferating. Further evidence indicating induction of cell senescence following treatment with these lower doses of doxorubicin was obtained by combined EdU labeling and CellMask Blue staining, and by staining the cells for β-galactosidase (Fig. 1D, E). In contrast, treatment with 10 µM doxorubicin resulted in profound cell death, with nearly all cells eliminated by day 4 of the time course.

To further quantify the cellular responses to varying doses of doxorubicin in a manner appropriate for distinguishing the behavior of distinct sub-populations of cells, we used quantitative live cell imaging (Incucyte™) to measure cell proliferation, and flow cytometry to monitor apoptotic and cell cycle responses at the single cell level. Flow cytometry measurements of apoptosis were performed by immunostaining the cells for simultaneous activation of the executioner caspase, caspase-3, and cleavage of PARP, a caspase-3 substrate (Janes et al., 2005) (Fig. 2A). DMSO and 0.5 µM doxorubicin treatments did not induce apoptotic cell death, while 2 µM doxorubicin induced a small fraction of U2OS cells to undergo apoptosis at early times after treatment, but this early burst of apoptosis subsided by 72 hrs after treatment (Fig. 2Bi and S1). In contrast, 10 µM doxorubicin treatment induced a significant fraction of cells to apoptose over the entire four day time course with a biphasic response, resulting in the complete absence of proliferation and the death of nearly all cells by the conclusion of the experiment (Fig. 2Bi-ii). The cell number of DMSO treated cells continued to increase over the 4-day time course, consistent with ongoing proliferation, which was eliminated by treatment with as little as 0.5 µM doxorubicin (Fig. 2Bii).

**Figure 2.**
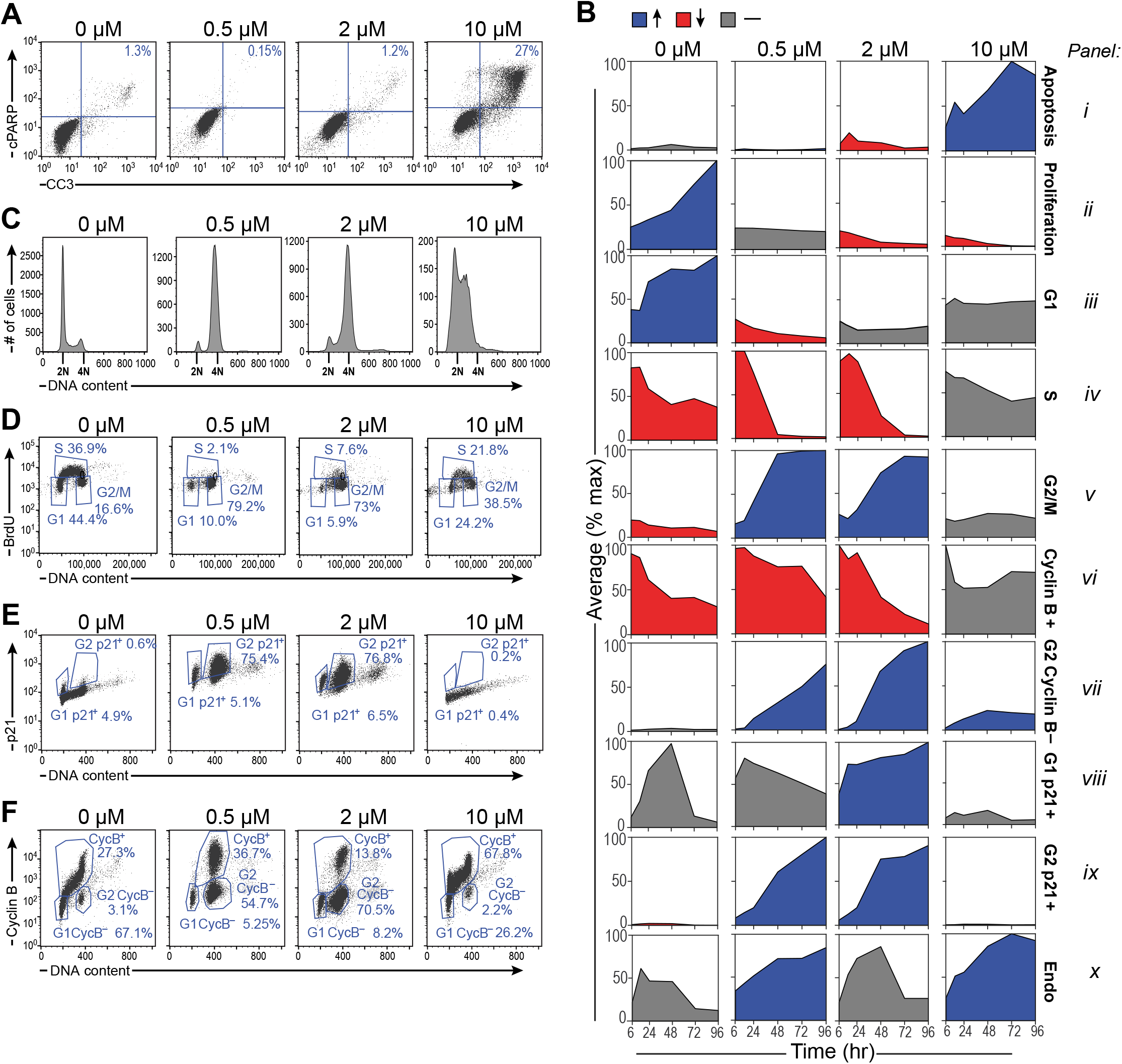
DNA damage induces G2/M arrest with dose-dependent differences between senescent p21^hi^/cyclin B^lo^ cells and apoptotic p21^lo^ cells. **A)** Representative flow cytometry scatter plot measuring apoptotic cells with cleaved caspase-3 (CC3) and cleaved PARP (cPARP) double positivity in U2OS cells 48 hours after doxorubicin treatment. **B)** Summary plots of mean phenotypic values (normalized to the maximum value across time and drug treatments) vs. time. Blue indicates a measurement that increases over time, red indicates a measurement that decreases, and gray indicates a measurement that remains the same. Tick marks represent 6, 24, 48, 72, and 96 hours after doxorubicin treatment. For boxplots of raw response data overlaid with replicate values, see supplemental figure S1. **C)** Histograms of DNA content in U2OS cells stained with propidium iodide (PI) 48 hours after doxorubicin treatment. 2N and 4N DNA content are annotated on the x-axis of the histogram plots. **D)** Representative flow cytometry scatter plots of BrdU antibody staining vs. PI staining in U2OS cells 48 hours after doxorubicin treatment. **E)** Representative flow cytometry scatter plot of p21 antibody staining vs PI staining in U2OS cells 96 hours after doxorubicin treatment. **F)** Representative flow cytometry scatter plot of cyclin B antibody staining vs PI staining in U2OS cells 96 hours after doxorubicin treatment.

Flow cytometry of propidium iodide-stained cells was used to monitor progression of U2OS cells through the cell cycle following treatment with doxorubicin (Fig. 2C). DNA replication activity was independently measured by analyzing the fraction of cells that incorporate BrdU in a 4 hour pulse (Fig. 2D). Mock-treated cells continue to proliferate, with cells distributed through all phases of the cell cycle. Cells treated with 0.5 µM and 2 µM doxorubicin proceeded through S phase and arrested with ∼80% or more of the cells in G2/M and the remaining cells in G1 (Fig. 2C, D), which developed over the course of 24 to 48 hours (Fig. 2Biii-v and S1). In contrast, under these treatment conditions, cells treated with 10 µM doxorubicin did not develop a pronounced G2 arrest, but instead entered and remained arrested in S phase, where they incorporated only low levels of BrdU and did not progress to G2 (Fig. 2Biv-v, 2D). This finding is consistent with our previous work indicating that this type of pulse doxorubicin treatment induced U2OS cells to undergo apoptosis in early S phase (Tentner et al., 2012). Cells were then stained for the presence or absence of Cyclin B and the CDK inhibitor p21^Waf1^ (Fig. 2Bvi-ix, 2E, and 2F). Consistent with the observed G2 arrest, 0.5 and 2 µM dox treatments caused marked up-regulation of the p21^Waf1^, while the apoptosis-inducing 10 µM dose did not (Fig. 2Bix, E). Cells arrested in G2 initially possessed high levels of cyclin B as would be expected, however, concomitant with the increase in p21, the level of cyclin B in these cells decreased to that of G1 cells (Fig. 2Bvi-vii, ix, 2F, and S1), indicating that low and intermediate doses of doxorubicin caused the cells to gradually lose the ability to transition from G2 to M phase. Taken together, these data indicate that 0.5 µM and 2 µM doxorubicin treatments arrest proliferation by inducing a G2 arrest that prevents cells from progressing into mitosis. Similarly to other cancer cell lines in which p16 has been silenced by methylation (Z. A. Stewart, Leach, & Pietenpol, 1999; Toettcher et al., 2009), a fraction of U2OS cells endoreduplicated in response to DNA damage as indicated by uptake of BrdU in cells containing >4N DNA (Fig. 2Bx, D).

To quantify the cell signaling response to different levels of DNA damage, 27 total measurements including the relative protein levels, protein phosphorylation, sub-cellular localization within the nuclear and cytoplasmic compartments, and heterogeneity between cellular sub-populations, for 19 signaling proteins representing key regulatory network nodes for cell cycle control, apoptosis, DNA damage response (DDR), and stress response signaling events (Fig. 3A) were quantified in 96-well plates using singleplex immunofluorescence microscopy at 6 time points: 6, 12, 24, 48, 72 and 96 hours after doxorubicin treatments. Representative signaling data at the 48 hour time point are shown in Figure 3B, with the complete quantified time courses shown in Figures 3C, S2 and S3. As expected, treatment of U2OS cells with doxorubicin strongly activated the DNA damage response, including up-regulation of γH2AX, stabilization of p53, and phosphorylation of the effector kinase Chk2 (Fig. 3B, C and S3). Interestingly, the DDIS-inducing doses of doxorubicin (0.5 and 2 µM) increased the level of p53 significantly more than the apoptosis-inducing dose (10 µM). In addition, these DDIS-inducing doses induced a large increase in the level of the cyclin dependent kinase inhibitor p21^Waf1^, which likely contributes to the U2OS cell cycle arrest, given the silencing by methylation of the p16 gene in this cell line (Fig. 3B, C) (Park et al., 2002).

**Figure 3.**
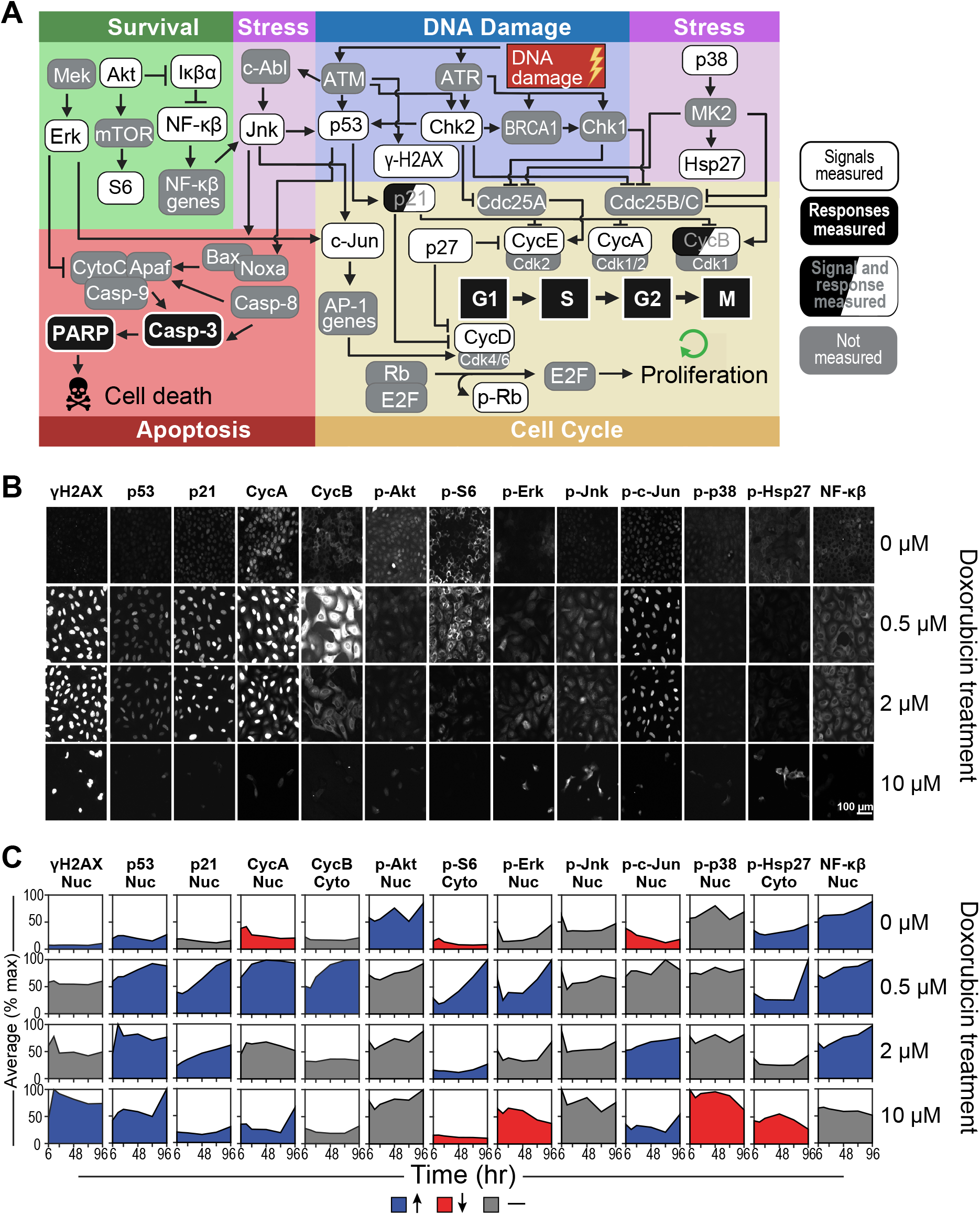
Quantitative single cell measurements define the DNA damage signaling landscape. **A)** A wiring diagram of the signaling network that modulates cell-cycle progression and apoptosis after DNA damage, created with BioRender.com. **B)** Representative immunofluorescence images of a subset of signaling proteins measured 48 hours after doxorubicin treatment. Each field of view represents a distinct well on a 96-well plate that was immunostained for 1 total or phosphoprotein. **C)** Quantification of the mean fluorescence intensity (normalized to the maximum value across time and drug treatments) over time in either the nuclear or cytoplasmic compartment, depending on the given protein measured. Blue indicates a measurement that increases over time, red indicates a decreasing measurement, and gray indicates a measurement that has remained the same. Tick marks on the x-axis represent 6, 24, 48, 72, and 96 hrs. For boxplots of raw signal data overlaid with replicate values, see supplemental figure S2.

Cell cycle arrest is a fundamental component of the DNA damage response. Consequently, we monitored the levels and localization of cyclins A, B, D and E, as well as phosphorylation of the retinoblastoma protein (Rb) (Fig. 3B, C and S3). In DMSO treated cells, the levels of these cell cycle regulating proteins remained largely unchanged for the 4 day duration of the experiment. The G2 arrest of the cells treated with 0.5 µM and 2 µM doxorubicin was reflected in the accumulation of cyclins A and B. Interestingly, U2OS cells undergoing DDIS accumulate cyclin E, indicating that they may be primed for another round of DNA replication. This is consistent with our previous data that HCT116 cells, which like U2OS cells have wild-type p53 and lack p16, also arrest in G2 after DNA damage and accumulate cyclin E, putting them in a 4N pseudo-G1 state that likely contributes to their propensity for endoreduplication (Toettcher et al., 2009). A small percentage (3-4%) of U2OS cells also endoreduplicated after DNA damage as shown by a population of 8N cells after the 0.5 and 2 µM doxorubicin treatments (Fig. S1).

Since signal transduction pathways other than the canonical DDR signaling network also contribute to cell fate determination, we measured the post-translational modifications and proteins levels reflective of Akt, Erk, JNK, p38 and NF-κB activity (Fig. 3B, C and S3). These pathways exhibit complicated, time- and doxorubicin-dose dependent behaviors that required data-driven modeling to integrate with the canonical DDR signaling network and downstream cell-fate responses (Janes & Yaffe, 2006).

### A Tensor PLSR Model Distinguishes Alternative Cell Fates after DNA Damage

To relate these complex time- and dose-dependent changes in cellular signaling to the observed phenotypic responses, we used tensor partial least square regression (t-PLSR) to identify signals, responses, and time points that significantly correlate with specific cell fates. Traditional “unfolded” PLSR is a widely used dimensional reduction modeling method in which the relationship between measured signaling events and phenotypic responses is inferred from maximizing the covariance between the two (Geladi & Kowalski, 1986; Janes et al., 2005). Independent signals, dependent cellular responses, and timepoints are weighed separately in the PLSR matrix formulation. In contrast, t-PLSR specifically preserves the natural structure of the data, regressing the stimulus-timepoint-response/cell fate tensor on the stimulus-timepoint-signaling tensor, linking these tensors via regression coefficients (Bro, 1996; Chitforoushzadeh et al., 2016). In addition to providing insight into how particular aspects of temporally evolving signaling activities are important for making predictions, tensor PLSR models use fewer parameters than unfolded PLSR, resulting in less overfitting, which is of particular importance when modeling a small number of treatments, as in our case.

In t-PLSR, the signal (X) and response (Y) tensors are simultaneously decomposed into three distinct individual matrices for each tensor: treatment scores (s_x_ or s_y_), signal (w_sx_) or response (w_sy_) weights, and time weights (w_tx_ or w_ty_) (Fig. 4A). To generate these matrices, the signaling tensor X and the cellular response tensor Y are jointly factored using a linear relationship between s_x_ and s_y_ in which the regression coefficients matrix is the slope within this relation, and this factorization occurs iteratively until the covariance between X and Y has been maximized. Upon convergence, these values are considered the score and weight values for latent variable #1, which is conceptually analogous to principal component 1 in traditional unfolded PLSR. The residuals are then subtracted from X and Y to compose the tensors used for the next round of factorization, which will be used to compute latent variable #2, and so on until a majority of the variance in the data has been explained. Treatment scores describe where certain treatments fall in the latent variable space, while signal and response weights describe the contribution of each individual signal or response to a specific latent variable. Time weights offer additional insight into which timepoints are weighted more heavily across all signals and responses in constructing a specific latent variable, information that is difficult to parse in traditional unfolded PLSR. As shown in figure 4A, the appropriate product of these scores and weights, when added over all latent variables, recapitulates the original signaling or response tensor.

**Figure 4.**
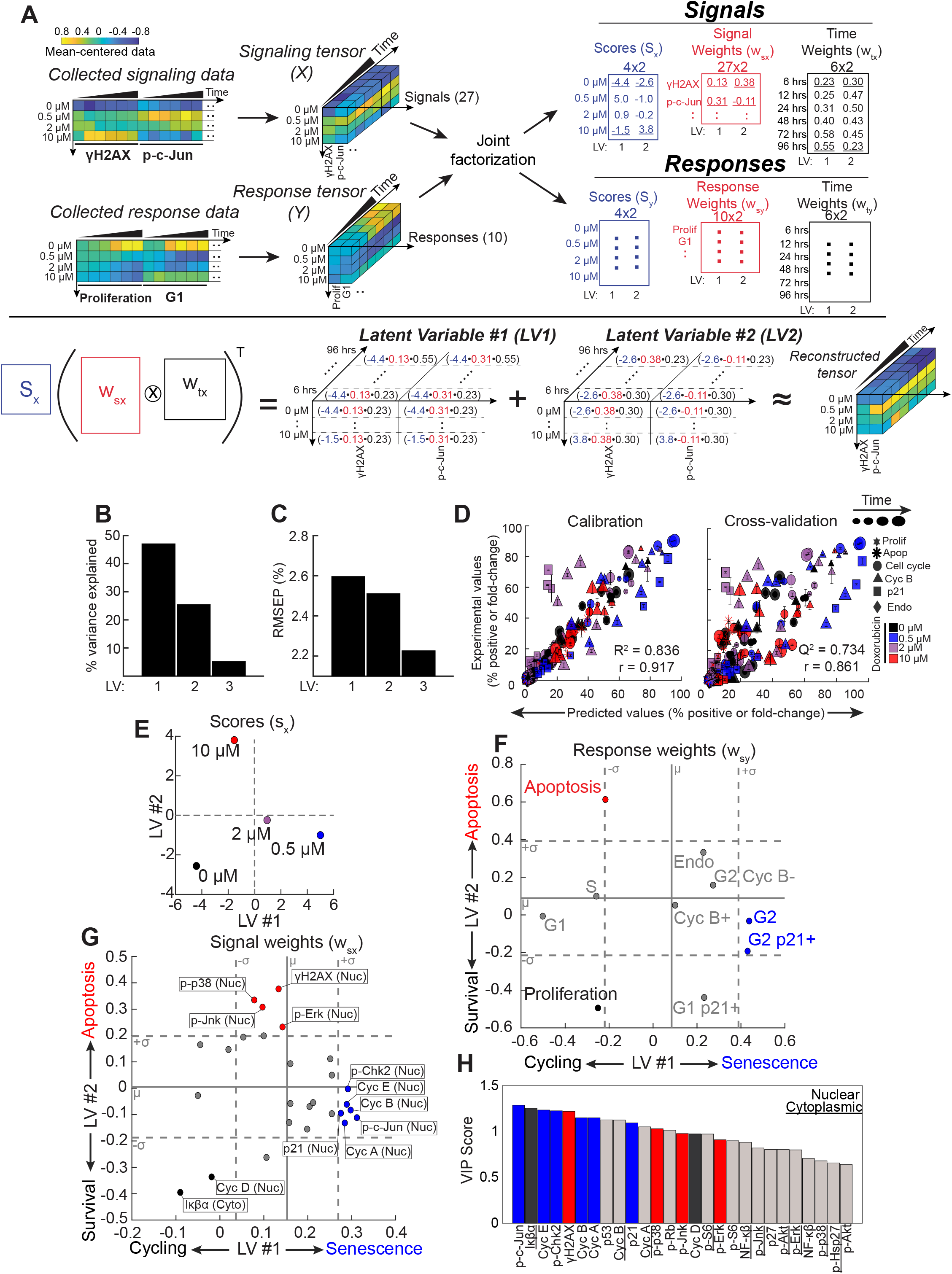
A Tensor PLSR model identifies latent variables that define a survival-apoptosis axis, and a cycling-senescence axis. **A)** A schematic of the “tensor” PLSR (t-PLSR) algorithm. The transpose of Khatri-Rao product of the computed w_sx_ and w_tx_ multiplied by the computed s_x_ should be able to fairly recapitulate the original tensor, as illustrated. **B)** Bar plots of the percent variance explained by each latent variables (LV). **C)** The root mean square of the prediction (RMSEP) of each LV. **D)** Experimental vs. predicted scatter plots for the values used for calibration (left) and those used for leave-one-out cross-validation (right). Error bars represent standard error of the mean (SEM). Computed R^2^, Q^2^, and Pearson correlation are also shown. **E)** Treatment scores from the signal tensor visualized on a scatter plot of latent variable #2 vs. latent variable #1. **F)** Response weights from the model plotted on a scatter plot of latent variable #2 vs. latent variable #1. Solid gray line indicates the mean (µ) of 500 null models, while dotted gray lines indicate +/- 1 standard deviation (σ) from the average of null models (see text for details). **G)** Signal weights from the model plotted on a scatter plot of latent variable #2 vs. latent variable #1. Solid and dotted gray lines calculated as described in subpanel F. **H)** Bar plot of variable importance in projection (VIP) scores of signals used in the model. Underlining of the signal name indicates a cytoplasmic signal, while the absence of underlining indicates a nuclear signal.

A t-PLSR model containing three latent variables captured greater than 75% of the variance in the response data while also minimizing the root-mean square error of prediction (Fig. 4B and 4C), with over 72% of the variance explained by latent variables #1 (LV1) and 2 (LV2). There was good concordance between the predicted and experimentally observed phenotypic responses during both model calibration and cross-validation (Fig. 4D). The largest discrepancies between the experimental and predicted values were observed at the 2 µM doxorubicin dose, likely due in part to the heterogeneity of cell fate responses observed at this particular dose (∼25% cumulative apoptosis and ∼75% senescence) (Fig. S4). When examining different phenotypic responses, rather than specific drug doses, the model performed best at predicting the percentage of cells in G1, S and G2/M in both the calibration and cross-validation data with Q^2^ values greater than 0.65 (Fig. S5).

To gain insight into how these latent variables correlated with cell fates, the treatment scores and the response weights from the model were explored. Plotting the treatment scores on LV1 vs LV2 revealed that DMSO treatment fell in the LV 1 and 2 negative quadrant, while the treatment scores of both doses of doxorubicin that induced senescence (0.5 and 2 µM) were LV1 positive and LV2 negative. Notably treatment with the 2 µM dose, which induces a more heterogeneous mix of senescent and apoptotic cells, projected less positively on LV1 than did the 0.5 µM dose. In contrast, the treatment score of the apoptotic dose of doxorubicin (10 µM) projected negatively on the LV1 axis, but was strongly LV2 positive (Fig. 4E). Thus, treatments involving senescent doses of this genotoxic agent are distributed along LV1 while LV2 distinguished the apoptotic dose treatment from the rest.

To further refine the biological meaning of LV1 and 2, the cellular response weights were plotted, and the significance of their projections on these axes evaluated using statistical bootstrapping (Fig. 4F) (Caulk & Janes, 2019). 500 separate null models were constructed from randomly shuffled data, and the observed response weights corresponding to the real data then compared with those obtained from the null models. Responses whose weights were greater than one standard deviation from the mean of the null models were considered significant. Using this cut-off, proliferation emerged as significant within the negative LV1 and 2 quadrant, while the G2 and G2 p21+ state emerged as the only significant responses that were LV1 positive. This observation indicates that G2 arrested/sensencent cells are separated from cycling proliferative cells by progression along the LV1 axis, in excellent agreement with the distribution of senescence-inducing treatment scores along this axis (2 µM and 0.5 µM doxorubicin; Fig. 4E). In contrast, apoptosis emerged as the only response that was significantly positive along the LV2 axis, in excellent agreement with the observation that the treatment score for the apoptotic dose of doxorubicin projected strongly in the positive LV2 direction. Measurements of the G1- and S-phase cell populations were not significantly distributed along the LV2 axis, they were significant on the negative LV1 axis (Fig. 4F), consistent with the location of actively cycling cells. The apoptosis response was narrowly below the cut-off for significance on the LV1 axes, which is consistent with the observation that cells treated with the apoptotic dose of doxorubicin maintained steady levels of cells in G1 and S that were not dissimilar from cells treated with the DMSO vehicle control (Fig. 2Biii-iv). Taken together, these observations, paired with the treatment scores, indicate that LV1 reflects a cycling versus senescence axis, while LV2 reflects survival versus apoptosis.

### Tensor PLSR and PCA Identify Signaling Pathways that Dictate Cell Fate

Interrogation of where specific molecular signals fall on the LV1 and 2 axes can infer potential causal relationships between signaling pathways and cell fates. Significant contributions of each of the 27 signaling measurements to LV1 and 2 were therefore evaluated using statistical bootstrapping as described above (Fig. 4G), and the relative importance of significant contributing signals further assessed using Variable Importance in Projection (VIP) scores (Fig. 4H) (Favilla, Durante, Vigni, & Cocchi, 2013). Both nuclear cyclin D and cytoplasmic Iκβα levels were significant contributors to LV1, projecting negatively along this axis, correlating with cell proliferation (Fig. 4G). These two elements in the t-PLSR model are in strong agreement with our interpretation of the LV1 axis, since the G1 cyclin, Cyclin D, is well-established as a key regulator of cell proliferation (Baldin, Lukas, Marcote, Pagano, & Draetta, 1993; Yang, Hitomi, & Stacey, 2006), while activation and nuclear translocation of NF-κβ, which is inhibited by Iκβα, has been shown to be elevated in senescent cells (Chien et al., 2011; Rovillain et al., 2011; Tilstra et al., 2012).

Nuclear γH2AX, together with levels of activated p38MAPK, JNK, and Erk in the nucleus projected positively along LV2, correlating with apoptosis. These findings further support our biological interpretation of the LV2 axes, since γH2AX intensity reflects the extent of DNA damage, and we and others have shown previously that p38MAPK, Erk and JNK have complex, context-dependent roles in cellular stress and DNA damage responses (Manke et al., 2005; Spallarossa et al., 2010; Tentner et al., 2012; Wada et al., 2008; X. Wang, Martindale, & Holbrook, 2000; Xia, Dickens, Raingeaud, Davis, & Greenberg, 1995), with JNK commonly associated with certain types of stress-associated cell death (Chen, Meyer, & Tan, 1996; Tournier et al., 2000; Verheij et al., 1996; Zanke et al., 1996).

Finally, nuclear levels of phospho-Chk2, Cyclin E, Cyclin A, Cyclin B, and p21^Waf1^ projected strongly along LV1 in a statistically significant manner, correlating closely with cell senescence. Chk2 has been previously associated as a driver of replicative senescence (Gire, Roux, Wynford-Thomas, Brondello, & Dulic, 2004; Nayak et al., 2017). p21^Waf1^ is both a canonical marker of senescence and the CDK inhibitor likely contributing to cell cycle arrest in this system (Brown, Wei, & Sedivy, 1997; Noda, Ning, Venable, Pereira-Smith, & Smith, 1994; Y. Wang, Blandino, & Givol, 1999), rationalizing the observed high levels of Cyclins E, A, and B in these 4N G2-arrested p21+ cells (Fig. 4F). The most unanticipated finding, however, was the very strong contribution of nuclear phospho-c-Jun to LV1, where it emerged as the strongest correlate with cell senescence based on the projection of its weight on LV1 and its VIP score, ranking as a more important contributor than p21^Waf1^ (Fig. 4G and H). c-Jun is a major component of the AP-1 transcription factor family, and has been well-characterized as a modulator of cell proliferation, cell-cycle progression, and cell death in a context-dependent manner (Johnson, Van Lingen, Papaioannou, & Spiegelman, 1993; Kovary & Bravo, 1991; Le-Niculescu et al., 1999; Shaulian et al., 2000). However, its potential role in modulating DNA damage-induced cell senescence has not been explored.

Phosphorylation and nuclear translocation of c-Jun is known to be mediated by JNK, Erk, and p38MAPK (Dérijard et al., 1994; Humar et al., 2007; Leppä, Saffrich, Ansorge, & Bohmann, 1998; Pulverer, Kyriakis, Avruch, Nikolakaki, & Woodgett, 1991), suggesting that the time-dependent activity of one or more of these kinases might be involved in regulating DDIS. Because time weights in t-PLSR reflect the aggregate measurements contributed by all signals or responses at one particular point in time, and are not amenable for dissecting the specific time-dependent contributions of any individual molecular signal, we instead applied principal component analysis (PCA) to the signaling data to parse how individual signals co-varied and separated by time point. Two principal components (PCs) captured greater than 90% of the variance in the signaling data. When each of the doxorubicin treatments was scored along these signal-generated PCs, the senescence-inducing doses (0.5 and 2 µM) segregated along PC1, while the apoptotic dose (10 µM) separated along PC2 (Fig 5A), essentially recapitulating the same proliferation-senescence and survival-apoptosis axes as those obtained using t-PLSR. This secondary analysis suggests that the largest variations in time-dependent signaling were informative for the doxorubicin-induced cell phenotypes in the dataset.

**Figure 5.**
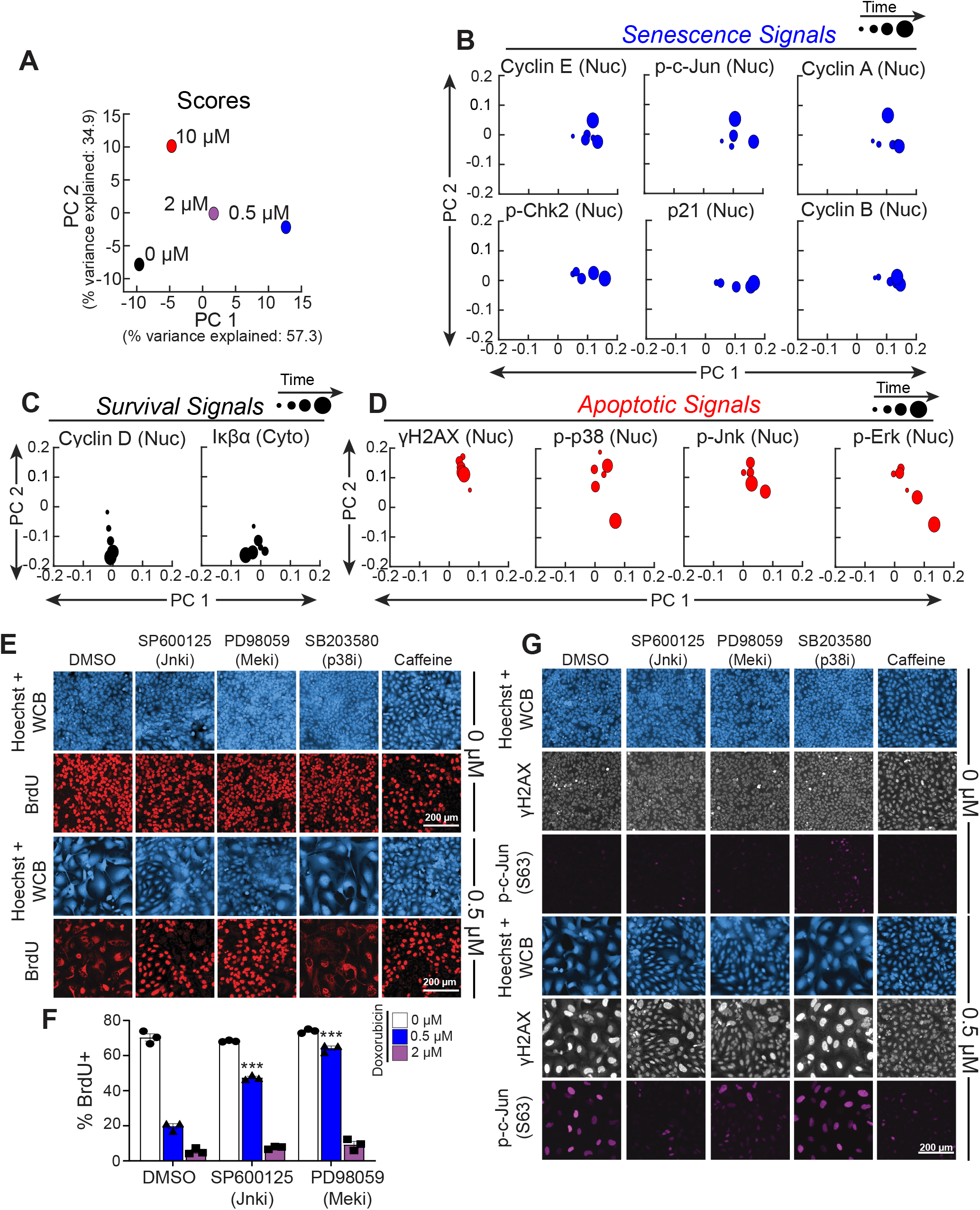
Kinases upstream of c-Jun, JNK and Erk, regulate senescence after DNA damage. **A)** PCA scores of doxorubicin treatments plotted on a scatterplot of principal component 2 (PC 2) vs. principal component 1 (PC 1). **B-D)** PCA loadings of significant **B)** senescent signals, **C)** survival signals, and **D)** apoptotic signals as determined by tensor PLSR. Larger points correspond to later timepoints. **E)** Representative immunofluorescence of cells treated with doxorubicin and either DMSO, 10 µM SP600125, 10 µM PD98059, 10 µM SB203580, or 5 mM caffeine. Doxorubicin was left on for 4 hours and then washed off, while inhibitors were left on the entire duration of the six-day experiment. Cell morphology was visualized with Hoechst staining and Whole Cell Blue (WCB) dye, and proliferative cells were visualized with nuclear BrdU antibody staining. **F)** Quantification of the percent of cells with that are positive for nuclear BrdU in the DMSO, 10 µM SP600125, and 10 µM PD98059 co-treatment conditions 6 days after doxorubicin treatment. Bars represent mean of three biological replicates, error bars represent SEM. ***: p < 0.001 with a two-way ANOVA and post-hoc Dunnett’s test vs. DMSO inhibitor control at the same dose of doxorubicin. **G)** Representative immunofluorescence for p-c-Jun(Ser63) and γH2AX of cells treated in the same conditions as in subpanel E.

PCA loadings of each of the molecular signals previously determined to be statistically significant contributors to LV1 and 2 by t-PLSR were then examined for their contributions to PC1 and PC2. Signals previously associated with senescence by t-PLSR were found, in PCA analysis, to become more PC1 positive over time (Fig. 5B), while signals associated with cell survival in t-PLSR became more PC2 negative over time in the PCA plots (Fig. 5C), consistent with PC1 representing proliferation-senescence and PC2 representing survival-apoptosis. Signals associated with DNA damage-induced apoptosis in t-PLSR, however, were more complex (Fig. 5D). All of the γH2AX signals remained clustered tightly in the positive PC2 direction, however signals for activated nuclear p38MAPK, JNK and Erk, while initially localized along the positive PC2 axis, were noted to become progressively less PC2 positive and more PC1 positive over time. These observations suggest potential nuanced and complex time-dependent roles for these three kinases after DNA damage, with potential roles in both apoptosis and DDIS.

### Early, but not Late JNK and Erk Signaling Controls Senescence After Low-Dose Doxorubicin Treatment

To experimentally investigate whether p38MAPK, Erk, and JNK directly contribute to regulating DNA damage induced senescence through phosphorylation of c-Jun, U2OS cells were treated with doxorubicin and the kinase activities of p38, Mek1, and JNK blocked with the specific small molecule inhibitors SB203580, PD98059, and SP600125, respectively, and validated by Western blotting (Fig. S6). In these experiments, U2OS cells were treated with a 4 hour doxorubicin pulse, and given fresh media on days 1 and 3 post-damage. Kinase inhibitors were applied simultaneously with doxorubicin and maintained throughout the 6 day duration of the experiment. Proliferation and senescence were evaluated by measuring the extent of BrdU incorporation over 24 hours starting 5 days after doxorubicin treatment. As observed previously, DMSO-treated U2OS cells showed strong BrdU uptake in spite of reaching high cell densities, while 0.5 µM doxorubicin treatment alone drives DDIS as indicated by morphological changes and the lack of BrdU incorporation into nuclear DNA (Fig. 5E). Notably, inhibition of JNK using SP600125 markedly abrogated this DDIS response. In contrast, following treatment with 2 µM doxorubicin, the DDIS could not be reversed by SP600125 (Fig. 5E and 5F). Blocking Erk signaling by inhibiting Mek1 similarly abrogated the senescence response to 0.5 µM doxorubicin, but, as was the case for JNK inhibition, PD98059 did not abrogate 2 µM doxorubicin-induced senescence (Fig. 5E and 5F). The pharmacologic perturbations suggested a role for Erk and JNK in DDIS triggered by low-dose doxorubicin.

To simultaneously visualize DNA damage signaling and signaling through c-Jun on a cell-by-cell basis, cells were co-stained for the γH2AX and p-c-Jun(S63) (Fig. 5G). DMSO-treated cells exhibited a uniform, small size and low levels of γH2AX and phospho-c-Jun, while treatment with 0.5 µM doxorubicin induces a population of large, flat senescent cells that stained positively for γH2AX and phospho-c-Jun (Fig. 5G). Although some heterogeneity in the extent of phospho-c-Jun staining was noted in the senescent cells, treatment with either SP600125 or PD98059, which abrogated DDIS, markedly reduced γH2AX staining and nearly completely eliminated nuclear phospho-c-Jun staining.

It should be noted that the abrogation of 0.5 µM doxorubicin-induced senescence by either SP600125 or PD98059 was not complete. The sub-population of cells that escaped senescence and proliferated after exposure to 0.5 µM doxorubicin upon treatment with either SP600125 or PD98059 were characterized by “normal” U2OS morphology, incorporation of BrdU, and low levels of both γH2AX and phospho-c-Jun (Fig. 5G). In contrast, the residual sub-population of senescent cells that persisted demonstrated an enlarged “fried-egg” cellular morphology, lack of nuclear BrdU incorporation, and increased levels of both γH2AX and phospho-c-Jun, further supporting the correlation of phospho-c-Jun with induction of DDIS. Inhibition of p38MAPK with SB203580 failed to suppress DNA damage-induced senescence, nuclear γH2AX intensity, or phospho-c-Jun staining, while addition of 5 mM caffeine, which served as a positive control, abrogated all three, as would be expected from its ability to simultaneously inhibit all three of the DNA damage response kinases ATM, ATR and DNA-PK (Fig. 5E and G). Taken together, we interpret these data as evidence that signaling though JNK and Erk, but not p38MAPK, plays a causal role in the induction of DDIS, likely through the phosphorylation of c-Jun (see below).

The DNA damage-induced senescent state, which is characterized by morphological changes, elevated cyclins E, A, and B, stable G2 arrest, and high levels of p21 emerges 3-4 days after DNA damage (Fig. 1-3). This multi-day time course between DNA damage and the emergence of the senescence state suggests a series of dynamic temporally-regulated signaling events and regulatory transitions that coordinate progression to senescence. To investigate at what point after DNA damage JNK and Erk signaling control the senescence cell fate decision, small molecule inhibitors were added either during the first 12 hours after the onset of doxorubicin-induced DNA damage, or added 12 hours after doxorubicin treatment and removed at 24 hours (Fig. 6A). As shown in Figures 6B and C, addition of the JNK inhibitor SP600125, the Mek1 inhibitor PD98059, or caffeine, during the first 12 hours after DNA damage significantly reduced the fraction of cells that underwent DDIS in response to the 0.5 µM doxorubicin treatment, with a smaller but still significant inhibition of DDIS in response to the 2 µM treatment as well. The p38MAPK inhibitor SB203580 did not reverse the DDIS phenotype, consistent with the prior results (Fig. 5E). In contrast, if added 12 hours after genotoxic stress, the JNK and Mek inhibitors and caffeine were unable to abrogate the DDIS in U2OS cells (Fig. 6D and E). Taken together, these observations suggest that JNK, Erk and DNA damage signaling – but not p38 signaling – are required within the first 12 hrs after DNA damage to initiate DDIS after low dose doxorubicin treatment.

**Figure 6.**
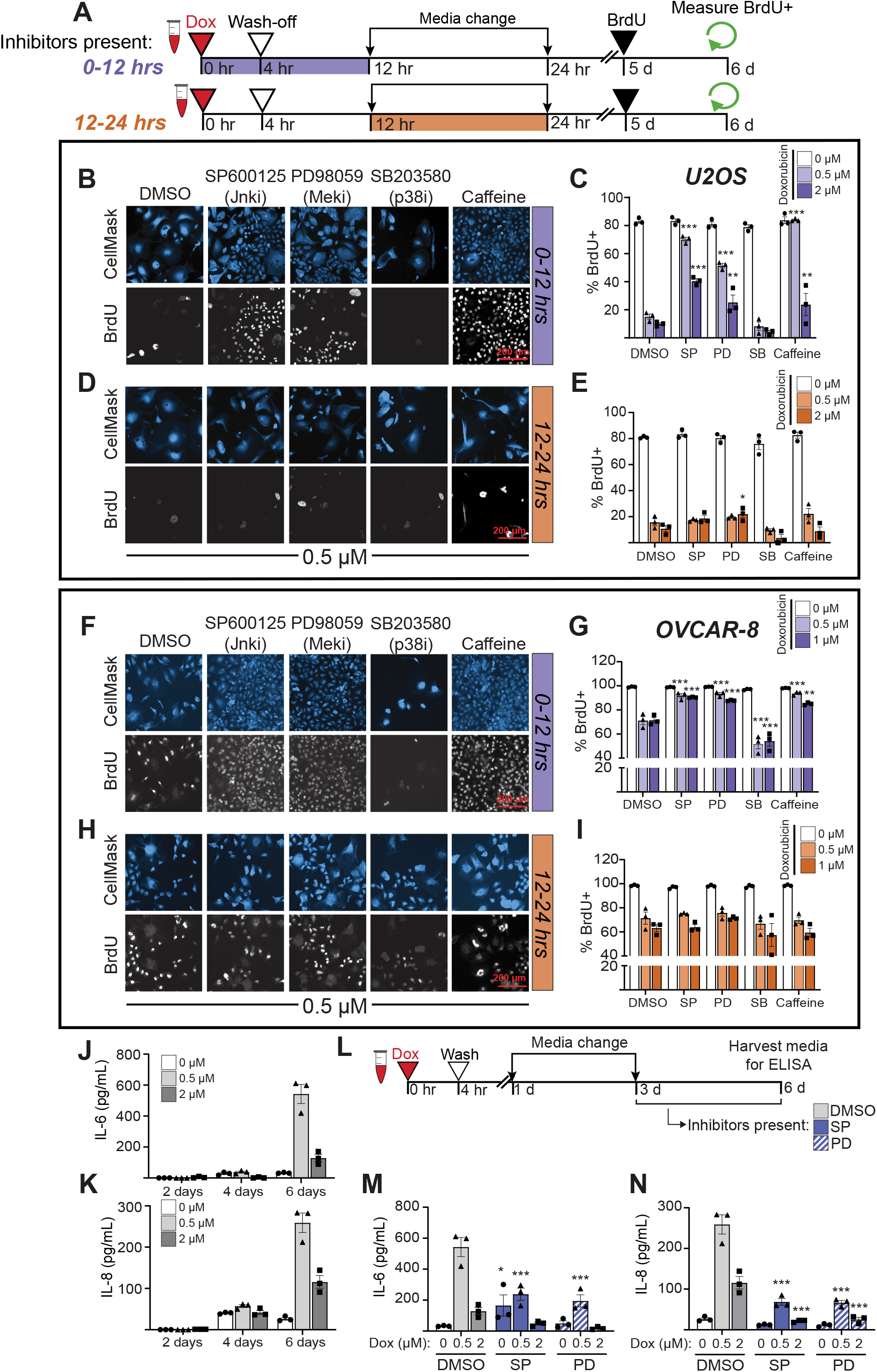
JNK and Erk signaling in the first 12 hours after DNA damage controls the senescence-proliferation decision, while late JNK and Erk signaling controls the SASP. **A)** Schematic of experimental workflow for U2OS and OVCAR-8 cells, using icons created in BioRender.com. **B)** Representative immunofluorescence (IF) of BrdU incorporation into DNA in U2OS cells treated with inhibitors (10 µM for SP600125, PD98059, and SB203580 drugs, and 5 mM of caffeine) for 0-12 hrs and co-stained with HCS CellMask Blue. **C)** Quantification of the percent of nuclear BrdU+ in U2OS cells after the 0-12 hour inhibitor condition. **D)** Representative IF of BrdU incorporation into DNA in U2OS cells treated with inhibitors for 12-24 hrs and co-stained with HCS CellMask Blue. **E)** Quantification of the percent of nuclear BrdU+ in U2OS cells after the 0-12 hour inhibitor condition. **F)** Representative IF of BrdU incorporation into DNA in OVCAR-8 cells treated with inhibitors (10 µM for SP600125, PD98059, and SB203580 drugs, and 1 mM of caffeine) for 0-12 hrs and co-stained with HCS CellMask Blue. **G)** Quantification of the percent of nuclear BrdU+ in OVCAR-8 cells after the 0-12 hour inhibitor condition. **H)** Representative IF of BrdU incorporation into DNA in OVCAR-8 cells treated with inhibitors for 12-24 hrs and co-stained with HCS CellMask Blue **I)** Quantification of the percent of nuclear BrdU+ in OVCAR-8 cells after the 12-24 hour inhibitor condition. **J)** IL-6 and **K)** IL-8 levels in culture media of U2OS cells 2, 4, and 6 days after doxorubicin treatment as measured by ELISA. **L)** Schematic of experimental workflow for ELISA-based co-inhibitor experiment **M)** IL-6 and **N)** IL-8 levels as measured by ELISA in U2OS tissue culture media 6 days after doxorubicin addition. DMSO inhibitor control bars in subpanels M and N are the same values as seen in subpanels J and K six days after doxorubicin. For all bar graphs, bars represent mean ± SEM of three biological replicates. ***: p < 0.001, **: p < 0.01 *: p < 0.05 with a two-way ANOVA and post-hoc Dunnett’s test for subpanels C, E, G, I, M, and L. All comparisons were vs. the DMSO treatment at the same dose of doxorubicin.

To examine whether the roles of JNK and Erk signaling in driving DDIS were unique to U2OS cells, similar studies were performed in OVCAR-8 high-grade serous ovarian cancer cells, a tumor type that is treated clinically with doxorubicin in the refractory or recurrent setting (Muggia et al., 1997; Yuan et al., 2021). Based on dose-response studies (Fig. S7), doxorubicin doses at or below 1 µM in this cell type caused little cell death, but instead resulted in the appearance of a sub-population of cells with enlarged nuclei and cytoplasm (i.e. a fried-egg appearance) that failed to incorporate nuclear BrdU by 6 days after treatment (Fig. 6F and G), albeit to a lesser extent than that seen in U2OS cells. Addition of SP600125, PD98059, or caffeine during the first 12 hours after DNA damage, resulted in disappearance of this subpopulation, with the emergence of a dense BrdU-positive monolayer of cells, indicating that JNK, Erk, and DDR inhibition early after damage can reverse the DDIS in this cell type (Fig. 6F and G), recapitulating the results obtained in U2OS cells. Similar to what was observed in U2OS cells, addition of JNK or Mek1 inhibitors, or treatment with caffeine, was unable to abrogate DDIS in OVCAR-8 cells if these were added later than 12 hours after genotoxic stress (Fig. 6H and I). These data indicate that, in both U2OS and OVCAR-8 cells, JNK and Erk signaling contribute to the early information processing that results in DDIS, and this commitment is made within 12 hours after genotoxic exposure.

### Late JNK and Erk activity contribute to the secretory associated secretory phenotype

Phospho-JNK and phospho-Erk levels are elevated at early timepoints, and the activity of these kinases influences the senescence/proliferation fate decision (Fig. 5 and 6A-I). However, their levels also remain elevated after low-dose doxorubicin at later timepoints (Fig. 3B), raising the question of whether there may be another function for these signaling pathways at late times. Since senescent cells are known to secrete pro-inflammatory cytokines, particularly IL-6 and IL-8, in response to oncogenic stress or DNA damage (Coppé et al., 2010, 2008; Kuilman et al., 2008; Rodier et al., 2009), and the JNK and Erk pathways are known to regulate cytokine secretion and mediate cytokine signaling in response to other non-DNA damage stimuli (Hoffmann et al., 2005; Krause et al., 1998), we hypothesized that the late phase JNK and Erk signaling after genotoxic stress might contribute to regulating the DDIS SASP.

To examine this, U2OS cells were treated with senescence-inducing doses of doxorubicin, and the levels of IL-6 and IL-8 assayed in the media at 2, 4, and 6 days after treatment. As shown in Fig. 6J and 6K, both 0.5 µM and 2 µM doxorubicin treatment induced IL-6 and IL-8 secretion that became significantly elevated by 6 days after treatment. Cells treated with 0.5 µM doxorubicin secreted significantly more IL-6 and IL-8 than cells treated with 2 µM doxorubicin, and the kinetics of cytokine release came after the senescence-associated morphology change. Addition of JNK or Mek1 inhibitors was therefore performed 72 hours after DNA damage to allow for the initiation of senescence, but prior to the detection of SASP-associated cytokines (Fig. 6L). As shown in Fig. 6M and N, addition of either the JNK inhibitor SP600125, or the Mek1 inhibitor PD98059 reduced the senescence-associated secretion of IL-6 and IL-8. Together with the data in panels A-I, this data indicates that JNK and Erk signaling regulate two different properties of cells undergoing DDIS on two distinct timescales, with early signaling implicated in the senescence decision, and late signaling involved in the senescence associated cytokine secretory response.

### JNK and Erk Signal Through AP-1 to Drive Cellular Senescence After Doxorubicin-Induced DNA Damage

Based on the findings in U2OS cells using t-PLSR and PCA analysis, we initially hypothesized that JNK, Erk, and p38MAPK acted through the phosphorylation of c-Jun to drive DDIS, and subsequently demonstrated through inhibition experiments that JNK and Erk activity, but not p38MAPK, were critical for the decision between senescence and proliferation within the first 12 hours after low-dose doxorubicin treatment. To demonstrate that c-Jun phosphorylation was directly regulated by these kinases during this early timeframe, Western blotting and immunofluorescence assays for phospho-c-Jun were performed at 6 and 12 hours following treatment of U2OS cells with 0.5 µM doxorubicin in the presence or absence of JNK or Mek1 inhibitors (Fig. S8 and S9). SP600125 caused a marked decrease in the levels of phospho-c-Jun after DNA damage in both assays, while PD98059 caused a more moderate, but also statistically significant reduction in c-Jun phosphorylation. Next, to examine whether inhibition of JNK or Mek1 activity in the presence of DNA damage controls the downstream transcriptional activity of c-Jun, we used an AP-1 mCherry-based transcriptional reporter containing three canonical AP-1 motifs within a minimal promoter upstream of mCherry. U2OS cells stably transfected with the AP-1 reporter were treated with DMSO or 0.5 µM doxorubicin for four hours, in the presence or absence of SP600125 or PD98059, and mCherry expression quantified by flow cytometry. As shown in Fig. 7A, doxorubicin treatment induced a ∼3 fold-increase in mCherry expression, similar to that observed in the PMA-treated positive control. Both the JNK inhibitor SP600125, and the Mek1 inhibitor PD98059, suppressed AP-1 transcriptional upregulation following DNA damage, mirroring the extent of suppression of c-Jun phosphorylation seen using these inhibitors by both immunoblotting and immunofluorescence (Fig. 7A, S8 and S9). To further validate that c-Jun is the relevant AP-1 family member in U2OS cells, the cells were transfected using a dominant negative c-Jun construct lacking the transactivation domain and sites of JNK and Erk phosphorylation, hereafter referred to as JunDN (Thompson, Gupta, Stratton, & Bowden, 2002). Expression of JunDN reduced AP-1-driven transcription in both the absence and presence of doxorubicin treatment, but importantly, reduced AP-1 transcription following doxorubicin treatment to the same level as that seen in untreated control cells (Fig. 7A).

**Figure 7.**
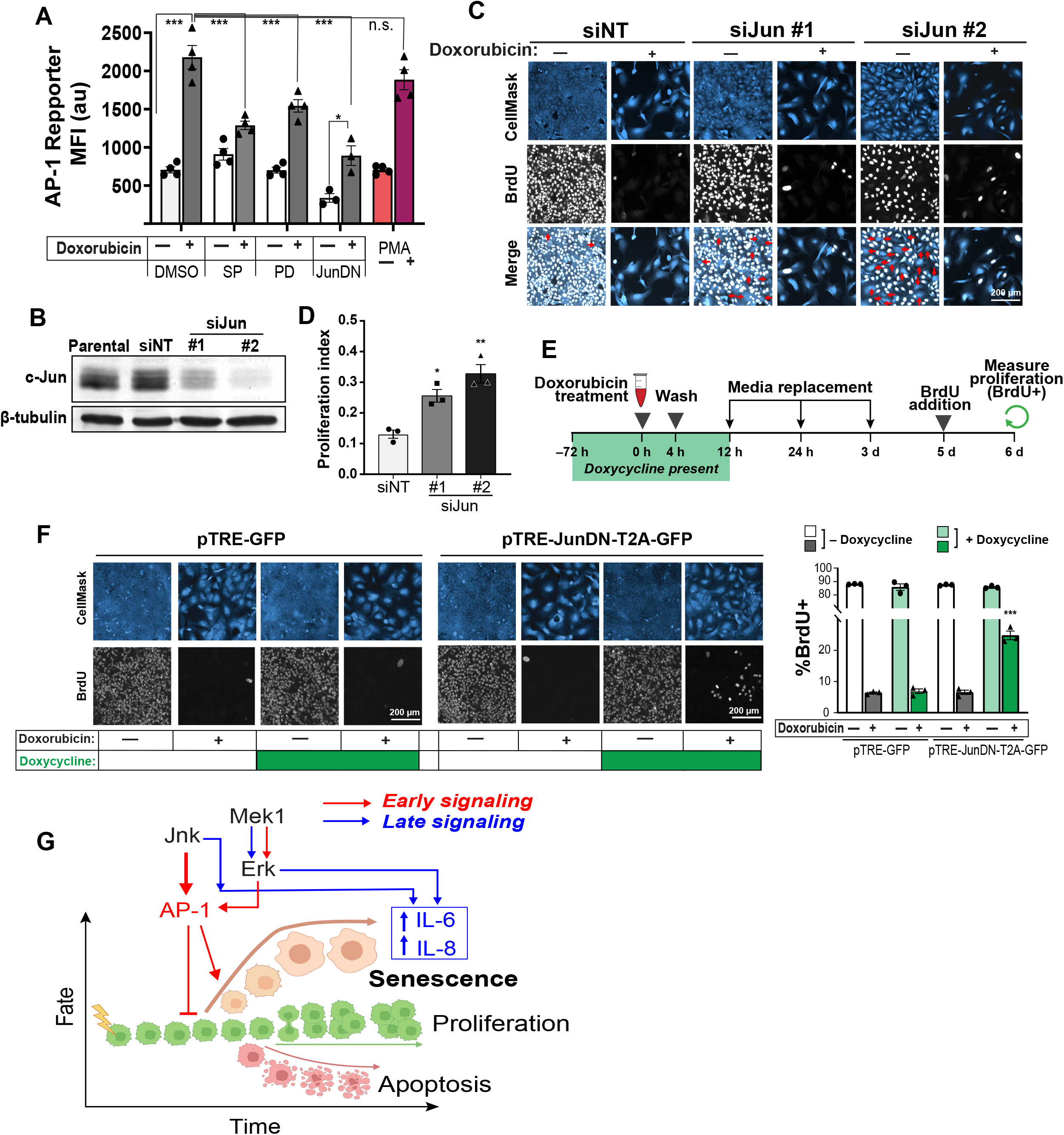
c-Jun directly controls the senescence-proliferation switch in U2OS cells after low doses of doxorubicin. **A)** Mean-fluorescence intensity (MFI) in the mCherry channel as measured by flow cytometry in U2OS cells expressing AP-1-mCherry reporter 24 hours after doxorubicin addition or continuous treatment with 100 nM (PMA). Bars represent mean ± SEM of biological replicates n = 4 (DMSO, SP, PD, and PMA) or n = 3 (JunDN). ***: p < 0.001, *: p < 0.05, n.s.: p > 0.05 with a two-way ANOVA and post-hoc Bonferroni’s multiple comparisons test. **B)** Representative immunoblot for c-Jun and β-tubulin 48 hours post-transfection. **C)** Representative immunofluorescence images of U2OS stained with anti-BrdU and HCS CellMask Blue six days after either 0.5 µM doxorubicin treatment (+) or DMSO (-). Red arrows denote BrdU negative cells. **D)** Quantified proliferation index, which is calculated as the percentage of BrdU-positive cells treated with 0.5 µM doxorubicin divided by the percentage of BrdU-positive cells after vehicle treatment (DMSO) for each respective siRNA. Bars represent the mean ± SEM of three biological replicates. **E)** Schematic of experimental workflow for inducible over-expression experiment, using icons created in BioRender.com. **F)** Representative immunofluorescence images of U2OS stably transfected with either the TRE-GFP or the TRE-JunDN-T2A-GFP stained with anti-BrdU and HCS CellMask Blue six days after doxorubicin treatment, with quantification of nuclear BrdU positivity six days after doxorubicin. Bars represent the mean ± SEM of biological replicates (n = 3). ***: p < 0.001 with a two-way ANOVA and post-hoc Dunnett’s test vs. the TRE-GFP condition with no doxycycline at the same dose of doxorubicin. **G)** Proposed mechanism of JNK and Erk signaling in DDIS, created in BioRender.com

To directly test whether c-Jun controls the early cell fate decision after DNA damage, U2OS cells were transfected with two independent siRNAs targeting c-Jun, resulting in good knockdown of c-Jun at the protein level (Fig. 7B). When proliferation was measured by BrdU incorporation six days after a 4 hour pulse of either vehicle or 0.5 µM doxorubicin, a significant increase in the percentage of BrdU positive cells was observed that correlated with extent of knockdown (Fig. 7C and 7D). However, even in the absence of doxorubicin treatment, c-Jun knockdown resulted in a sub-population of cells that were BrdU negative (Fig. 7D, merge panel), likely as a consequence of prolonged doubling times that exceeded the 24 hour BrdU pulse. To mitigate the impact of continual c-Jun suppression on basal cell proliferation, we next generated U2OS cells stably expressing a tetracycline-inducible (tet-on) construct containing JunDN linked with a T2A cleavable linker to GFP (JunDN-T2A-GFP). Expression of JunDN was induced by doxycycline for 3 days prior to a 4 hour drug pulse of 0.5 µM doxorubicin, and the cells then maintained in doxycycline for 12 hours after doxorubicin treatment. The media was then changed to doxycycline-free media, which was replaced with fresh doxycycline-free media 1 and 3 days after doxorubicin treatment (Fig. 7E). As shown in Figure 7F, control cells expressing a tet-on GFP-only construct exhibited no increase in BrdU positive cells six days after DNA damage, regardless of doxycycline treatment. In contrast, cells expressing the JunDN-T2A-GFP construct demonstrated a significant increase in the percentage of BrdU positive cells after DNA damage only in the presence of doxycycline (Fig. 7F and 7G). These data, combined with the c-Jun knockdown results in panels B-D confirms that AP-1 has a direct role in the senescence cell-fate decision after low-dose doxorubicin treatment, while the AP-1 reporter results, coupled with the phospho-c-Jun measurements in the presence or absence of inhibitors, indicate that JNK and Erk are the upstream kinases activating AP-1 in this context.

## DISCUSSION

In this manuscript, we interrogated canonical and non-canonical DNA damage signaling pathways for their influence on cell fate decisions in response to different levels of DNA damage. Since activation of a common set of DDR-activated signaling molecules, including among others ATM, Chk2, H2AX, Nbs1, 53BP1, p53 and p21 (Cuella-Martin et al., 2016; Fernando et al., 2011; Hsu, Altschuler, & Wu, 2019; Jackson & Bartek, 2009; Reyes et al., 2018) results in diverse phenotypic outcomes, we hypothesized that cross-talk from additional signaling pathways likely influenced the cell fate decision process. We were particularly interested in examining the signaling responses at the single cell level under conditions where subpopulations of cells underwent different cell fates using quantitative microscopy, immunofluorescence, and flow cytometry, combined with computational modeling. The resulting compendium of data was used to construct a t-PLSR model, using significantly fewer parameters to predict the responses than are required for a conventional unfolded PLSR models (Janes et al., 2005; Lee et al., 2012). Our results demonstrate that a relatively small number of treatments was sufficient to construct a model that was robust and provided biological insight. Using t-PLSR modeling and PCA analysis paired with subsequent experimental validation, we identified unexpected roles for the MAP kinases JNK and Erk in modulating the early fate decision between DDIS and proliferation in the setting of low-dose DNA damage through the common downstream target of c-Jun, as well as a later role for these kinases in controlling the SASP cytokines IL-6 and IL-8 in committed senescent cells.

Our finding of a central role for JNK in enhancing the cell fate choice of senescence after modest DNA damage was unexpected, since JNK is best known to function as a stress-responsive regulator of apoptotic and non-apoptotic cell death. JNK promotes intrinsic apoptotic cell death, both *in vitro* and *in vivo* (Hübner, Barrett, Flavell, & Davis, 2008; N. J. Kennedy et al., 2003; Tournier et al., 2000) through a variety of mechanisms, including direct phosphorylation and activation of pro-apoptotic BH3 proteins (Donovan, Becker, Konishi, & Bonni, 2002; Lei & Davis, 2003), inactivation of the central scaffolding molecule 14-3-3 (Sunayama, Tsuruta, Masuyama, & Gotoh, 2005; Tsuruta et al., 2004), and phosphorylation and activation of p53 and/or p73 to increase the transcription of the pro-apoptotic BH3 family member PUMA (Jones, Dickman, & Whitmarsh, 2007; Oleinik, Krupenko, & Krupenko, 2007). In addition, novel roles for JNK in promoting necroptosis, pyroptosis, ferroptosis and autophagic cell death have recently been observed (Dhanasekaran & Premkumar Reddy, 2017). JNK activity even prevents p53-mediated cell senescence in unperturbed MEFs in culture (Das et al., 2007). Our findings of a specific role for JNK in inducing senescence, rather than preventing senescence or promoting cell death, following modest levels of doxorubicin-induced DNA damage, suggest a very specific context-dependence in which JNK signaling controls the fate choice between senescence, proliferation, and death.

Conversely, the Erk MAP kinase pathway has typically be associated with enhancement of cell proliferation and differentiation, rather than senescence induction, through the direct Erk-dependent phosphorylation of a large number of transcription factors, including members of the TCF, ERG, ERF, PEA3, AP-1, EGR families as well as Runx2 and c-Myc themselves (Lavoie, Gagnon, & Therrien, 2020), resulting in their nuclear translocation, enhanced DNA binding, and transcription of immediate early genes and G1 cyclins. In addition, Erk phosphorylation of the stem cell transcription factors Oct4, Klf4, and Klf2 has been shown to decrease their stability and thus lead to loss of pluripotency (M. O. Kim et al., 2012; Spelat, Ferro, & Curcio, 2012; Yeo et al., 2014). In the setting of DNA damage, our lab and others have shown previously that Erk contributes to both G1/S arrest and subsequent apoptotic cell death after DNA damage using high doses of doxorubicin or cisplatin (Tentner et al., 2012; X. Wang et al., 2000; Yeh et al., 2004). To our knowledge, a clear role for Erk in contributing to cell senescence following extrinsic DNA damage induced by low-dose cytotoxic chemotherapy has not been reported. Erk activity has, however, been shown to contribute to the induction of senescence in p53 wild-type cells in response to expression of oncogenic Ras and Raf (Lin et al., 1998; W. Wang et al., 2002; Zhu, Woods, McMahon, & Bishop, 1998). Interpretation of our findings in light of the findings from Bartek, Lucas, and colleagues, who showed that oncogene activation results in replication stress and intrinsic DNA damage, suggests that a similar Erk-dependent senescence pathway as that observed following oncogene activation is being activated by extrinsic DNA damage caused by low doses of genotoxic drugs (Bartkova et al., 2006).

Unexpectedly, we did not detect a quantitatively significant role for the p38MAPK pathway in controlling the senescence decision at early times following low-dose DNA damage, despite the fact that this pathway is known to be activated by many different types of stress, including DNA damage (Roux & Blenis, 2004). In response to more extensive DNA damage induced by higher doses of doxorubicin (or cisplatin), p38MAPK, acting through MAPKAP Kinase-2, is known to be required for sustained cell cycle arrest and cell survival, but this effect results from signaling at late times greater than 12 hours, and is important only in cells in which p53 activity is at least partially defective (Cannell et al., 2015; Morandell et al., 2013; Reinhardt et al., 2007, 2010). This delayed temporal activation of the p38MAPK-MK2 pathway following DNA damage (Reinhardt et al., 2010) rationalizes the lack of effect of p38MAPK inhibition on low dose DNA damage-induced senescence onset, and instead indicates roles for this pathway primarily at later times. Consistent with this, a major role for p38MAPK in regulating the SASP has been reported by Campisi and colleagues (Freund et al., 2011).

The AP-1 transcription factor in general, and c-Jun in particular, emerged as key determinants of senescence in our experiments. The AP-1 transcription factor is a hetero- or homo-dimeric complex comprised of members of the Jun, Fos, ATF, and MAF protein families, that plays an important role in oncogenesis and tumor proliferation, and is known to regulated by JNK, Erk, and p38MAPK in a context-specific manner (Eferl & Wagner, 2003). The major AP-1 family member that emerged from our experimental findings and computational t-PLSR and PC analyses of low-dose DNA damage in U2OS cells was c-Jun, whose phosphorylation and nuclear accumulation correlated with the early cell fate decision towards senescence. Suppression of c-Jun using siRNA partially reversed the senescence phenotype after DNA damage, and this partial bypass of senescence was further confirmed using inducible expression of a non-phosphorylatable dominant negative form of c-Jun, which suppressed AP-1 activity in the cells.

Intriguingly, two recent papers have implicated a role for AP-1 in oncogene induced senescence. Martinez-Zamudio *et al*. (Martínez-Zamudio et al., 2020) used a combination of transcriptomic, ATAC-Seq and ChIP-Seq data to nominate AP-1 as a pioneering transcription factor that altered chromatin structure and allowed the establishment of a senescence-inducing enhancer landscape following the inducible expression of a Ras^G12V^ mutant oncogene in WI-38 human fibroblasts. siRNA knock-down of c-Jun had a larger effect on the Ras-induced senescence transcriptional program than did knock-down of the non-pioneering transcription factors ETS1 and RelA, primarily suppressing transcription of SASP-related genes, but also partially re-activating proliferation-associated genes. Those authors did not examine upstream regulatory kinases, or demonstrate reversion of a senescence cellular phenotype. Nonetheless, their extensive and comprehensive epigenetic and transcriptional analysis of oncogene-induced senescence is in good potential agreement with our findings of an important role for c-Jun in doxorubicin-induced senescence in U2OS cells. In a separate study, Han *et al*. used inducible expression of a mutant Ras^G12V^ oncogene in hTERT-immortalized BJ fibroblasts and observed upregulation of enhancer RNAs that were enriched for the binding motif of AP-1 (Han et al., 2018), suggesting an important role for AP-1-driven gene transcription in response to oncogenic stress. They then identified a specific AP-1 driven enhancer region controlling the expression of FOXF1as critical for the onset of oncogene-induced senescence. Taken together, these two studies support our findings that JNK and Erk modulate an AP-1-driven program of senescence, and suggest strong parallels between oncogene-induced senescence and senescence induced by extrinsic DNA damaging drugs. Interestingly, Davis and colleagues have shown, using a PTEN inactivation-dependent model of prostate cancer, that genetic elimination of JNK, or its upstream activators MKK4 and 7, results in marked enhancement of prostate tumor growth by suppressing the senescence response of prostate cells upon PTEN loss (Hübner et al., 2012). Those findings are in excellent agreement with our proposed model for an important role of JNK signaling in promoting senescence in response to DNA damage (Fig. 7G), caused either by oncogenic replication stress, or by low doses of extrinsic genotoxic drugs.

Heterogeneity in the behavior and response of cancer cells is now well established (Dagogo-Jack & Shaw, 2018; Melo, Vermeulen, Fessler, & Medema, 2013), and recent work indicates a similar heterogeneity is present within senescent cells (Cohn, Gasek, Kuchel, & Xu, 2022). In this regard, it is interesting that we observed heterogeneity in the proliferative response of U2OS and OVCAR-8 cells following JNK and Mek inhibition in response to treatment with 0.5 µM doxorubicin, with only a fraction of the inhibitor-treated cells escaping from senescence onset. The molecular basis for this heterogeneous response is unclear, and future work will be required to better define the underlying mechanisms, which represent a general challenge for predicting the clinically relevant biology of complex tumors during both their development and their response to treatment (Chapman et al., 2014; Kemper et al., 2015; Marusyk et al., 2014; Morrissy et al., 2016; Quintana et al., 2010; Zhao, Hemann, & Lauffenburger, 2014). Nonetheless, our finding of distinct early and late roles for JNK and Erk in senescence progression is of clear clinical utility, given the recent interest in the use of senolytics and SASP inhibitors for the treatment of cancer (Fielder et al., 2022; Short, Fielder, Miwa, & von Zglinicki, 2019; L. Wang, Lankhorst, & Bernards, 2022). For example, administering a JNK or Mek inhibitor days or weeks after chemotherapy could favorably lower IL-6 and IL-8 levels secreted by treatment-induced senescent cells, and thus decrease the IL-6/IL-8-mediated signaling events in the tumor microenvironment that favor cancer progression (Coppé et al., 2008; Goulet et al., 2019; Ruhland et al., 2016; Wu et al., 2019); conversely, administering these inhibitors at the same time as chemotherapy could drive cells towards proliferation rather than senescence due to the early cell fate decision-making role of JNK and Erk, resulting in tumor resistance to cytotoxic agents and hindering the efficacy of senolytic therapies later on. These findings of distinct temporal and context-dependent roles for JNK and Erk MAP kinases in the setting of extrinsic genotoxic stress, however, further illustrate the extraordinary complexity and plethora of cellular responses mediated by these highly conserved signaling cascades in controlling the DNA damage response.

## ACKNOWLEDGEMENTS

We gratefully acknowledge Bjorn Millard for assistance with microscopy, Brian Joughin for computational advice, Leslie Gaffney for assistance with figure design, and all members of the Yaffe lab for support and suggestions. We thank the Koch Institute’s Robert A. Swanson (1969) Biotechnology Center for technical support, specifically the Flow Cytometry Core Facility and the High Throughput Sciences Core, which is supported in part by the Koch Institute Support Grant P30-CA14051 from the National Cancer Institute. This work was supported by NIH grants R35-ES028374 and R01-GM104047 to M.B.Y., U54-CA112967 to M.B.Y and D.A.L., P50-GM68762 to P.K.S. and D.A.L., an NSF Graduate Research Fellowship 1122374 to T.S.N, the Ovarian Cancer Research Fund, the Charles and Marjorie Holloway Foundation, and the MIT Center for Precision Cancer Medicine.

## AUTHOR CONTRIBUTIONS

T.S.N, G.J.O, and M.B.Y. designed the experiments, analyzed the data, performed computational studies, and wrote the paper. T.S.N., and G.J.O. performed the experiments. A.R.T., K.A.J., and D.A.L. contributed to the computational analysis of the data and wrote the paper, and P.K.S. designed the experiments and analyzed the data.

## MATERIALS AND METHODS

### Cell culture

U-2 OS (U2OS) and HEK293T cells were purchased from the American Type Culture Collection (ATCC), and cultured in complete media consisting of Dulbecco’s Modified Eagle’s Medium (Corning, catalog # 10-013-CV) supplemented with 10% fetal bovine serum (FBS, VWR, catalog # 97068-085) and 1% penicillin-streptomycin (pen-strep, Gibco, catalog # 15070063). OVCAR-8 cells were a gift from Dr. Paula Hammond, and cultured in RPMI (Gibco, catalog # 11875093) supplemented with 10% FBS and 1% pen-strep. Cell cultures were maintained in a humidified incubator at 37°C with 5% CO_2_, and cells with less than 30 passages were used for experiments.

### Antibodies and chemicals

Doxorubicin hydrochloride (catalog # PHR1789), SP600125 (catalog # S5567), and caffeine (catalog # C0750) were purchased from Sigma-Aldrich. PD98059 (catalog # 9900) was purchased from Cell Signaling Technology. SB203580 (catalog # 1202) was purchased from Tocris. Doxycycline (catalog # 631311) was purchased from Clontech. The following antibodies were used for immunofluorescence measurements:

| **Antibody** | **Vendor** | **Catalog #** | **RRID** | **Dilution** |
| --- | --- | --- | --- | --- |
| γ-H2AX | Millipore | 05-636 | RRID:AB_309864 | 1:200 |
| p-Akt | Cell Signaling Technology | 4058 | RRID:AB_331168 | 1:200 |
| p-Erk | Cell Signaling Technology | 4377 | RRID:AB_331775 | 1:200 |
| p-JNK | Cell Signaling Technology | 9251 | RRID:AB_331659 | 1:100 |
| NF-κB | Santa Cruz | sc-8008 | RRID:AB_628017 | 1:50 |
| p-p38 | Cell Signaling Technology | 4631 | RRID:AB_331765 | 1:200 |
| p-S6 | Cell Signaling Technology | 4856 | RRID:AB_2181037 | 1:200 |
| p27 | Santa Cruz | sc-1641 | RRID:AB_628074 | 1:50 |
| p-Hsp27 | Cell Signaling Technology | 2401 | RRID:AB_331644 | 1:200 |
| p53 | Santa Cruz | sc-263 | RRID:AB_628084 | 1:50 |
| p-c-Jun(S73) | Cell Signaling Technology | 9164 | RRID:AB_330892 | 1:200 |
| p-c-Jun(S63) | Cell Signaling Technology | 9261 | RRID:AB_2130162 | 1:200 |
| p-Chk2 | Cell Signaling Technology | 2661 | RRID:AB_331479 | 1:200 |
| IκBα | BD Biosciences | 610691 | RRID:AB_398014 | 1:200 |
| Cyclin D | Santa Cruz | sc-20044 | RRID:AB_627346 | 1:50 |
| p-Rb | Cell Signaling Technology | 9308 | RRID:AB_331472 | 1:200 |
| Cyclin E | Santa Cruz | sc-247 | RRID:AB_627357 | 1:50 |
| Cyclin A | Santa Cruz | sc-751 | RRID:AB_631329 | 1:50 |
| p21 | Millipore | 05-345 | RRID:AB_309684 | 1:200 |
| Cyclin B | Santa Cruz | sc-752 | RRID:AB_2072134 | 1:50 |
| BrdU | BD Biosciences | 555627 | RRID:AB_395993 | 1:400 |
| BrdU | Cell Signaling Technology | 5292S | RRID:AB_10548898 | 1:1000 |

For immunoblots, the following antibodies were used:

| **Antibody** | **Vendor** | **Catalog #** | **RRID** | **Dilution** |
| --- | --- | --- | --- | --- |
| p-c-Jun(S73) | Cell Signaling Technology | 3270 | RRID:AB_2129575 | 1:1000 |
| c-Jun | Cell Signaling Technology | 9165 | RRID:AB_2130165 | 1:1000 |
| p-Erk | Cell Signaling Technology | 4370 | RRID:AB_2315112 | 1:2000 |
| Erk | Cell Signaling Technology | 4696 | RRID:AB_390780 | 1:2000 |
| p-JNK | Cell Signaling Technology | 4668 | RRID:AB_823588 | 1:1000 |
| JNK | Cell Signaling Technology | 9252 | RRID:AB_2250373 | 1:1000 |
| p-MK2 | Cell Signaling Technology | 3007 | RRID:AB_490936 | 1:1000 |
| MK2 | Cell Signaling Technology | 3042 | RRID:AB_10694238 | 1:1000 |
| p-p38 | Cell Signaling Technology | 4631 | RRID:AB_331765 | 1:1000 |
| p38 | Cell Signaling Technology | 9212 | RRID:AB_330713 | 1:1000 |
| β-tubulin | Sigma-Aldrich | T7816 | RRID:AB_261770 | 1:2000 |
| GAPDH | Cell Signaling Technology | 97166 | RRID:AB_2756824 | 1:1000 |

For flow cytometry, the following antibodies were used:

| **Antibody** | **Vendor** | **Catalog #** | **RRID** | **Dilution** |
| --- | --- | --- | --- | --- |
| Cleaved Caspase-3 | BD Biosciences | 559565 | RRID:AB_397274 | 1:200 |
| Cleaved PARP | BD Biosciences | 558710 | RRID:AB_1645431 | 1:200 |
| Cyclin B | Santa Cruz | sc-752 | RRID:AB_2072134 | 1:50 |
| p21 | Millipore | 05-345 | RRID:AB_309684 | 1:200 |
| BrdU | BD Biosciences | 555627 | RRID:AB_395993 | 1:200 |
| Pacific Blue cleaved caspase-3 (OVCAR-8 data) | Cell Signaling Technology | 8788S | RRID:AB_2797665 | 1:50 |
| Alexa fluor 647 cleaved PARP (OVCAR-8 data) | Cell Signaling Technology | 6987S | RRID:AB_10858215 | 1:50 |

The following secondary antibodies were used for both immunofluorescence and flow cytometry in this study:

| **Antibody** | **Vendor** | **Catalog #** | **RRID** | **Dilution** |
| --- | --- | --- | --- | --- |
| Alexa Fluor 488 goat anti-rabbit | Invitrogen | A-11008 | RRID:AB_143165 | 1:200 (CC3)-2000 (BrdU, CycB) for FC, 1:1000 for IF |
| Alexa Fluor 647 goat anti-rabbit | Invitrogen | A-21245 | RRID:AB_141775 | 1:1000 for IF |
| Alexa Fluor 647 goat anti-mouse | Invitrogen | A-21236 | RRID:AB_2535805 | 1:2000 for FC, 1:1000 for IF |

### Doxorubicin treatment for signaling and response measurements

Cells were plated at least 24 hours before treatment in complete culture media. Doxorubicin or DMSO (Sigma-Aldrich, catalog #D8418) was directly added to culture media, and media was then removed 4 hours later and replaced with media containing 1% fetal bovine serum for the duration of the experiment for immunofluorescence, cell cycle, apoptosis, and proliferation assays.

### β-galactosidase activity assay

U2OS cells were plated in a 24-well plate 24 hours before doxorubicin treatment in complete media. Doxorubicin was directly added to the culture media, and 4 hours later the media was replaced with media containing 1% FBS for the duration of the experiment. The plate was harvested 4 days after doxorubicin, and the Senescence β-Galactosidase Staining Kit (Cell Signaling Technology, catalog #9860) was used according to the manufacturer’s protocol. Images were taken on an EVOS fluorescence microscope (Thermo-Fisher) in the brightfield channel.

### Immunofluorescence for BrdU

In experiments where 5-bromo-2’-deoxyuridine (BrdU, Thermo-Fisher, catalog #B23151) incorporation into DNA was measured with immunofluorescence, cells were seeded in optical bottom 96-well plates (Thermo-Fisher, catalog # 165305) at the beginning of the experiment, and then were incubated in 10 µM BrdU for 24 hours before fixation. Cells were fixed with 4% paraformaldehyde (Electron Microscopy Sciences, catalog # 15710) for 15 minutes at room temperature, washed 3x with phosphate-buffered saline (PBS, GenClone, catalog # 25-507B), and permeabilized with PBS containing 0.1% Triton-X (Sigma-Aldrich, catalog # T9284) for 20 minutes at room temperature. Cells were tehen incubated with a primary antibody solution containing BrdU antibody diluted 1:400 or 1:1000 in PBS containing 5% goat serum (Abcam, catalog # ab7481) and 0.1% Triton-X for 1 hour at room temperature. Afterwards, cells were washed 3x with PBS and incubated with an anti-mouse antibody conjugated to Alexa Fluor 647 diluted 1:1000 in PBS for 1 hour at room temperature. Cells were then washed 3x with PBS, and incubated either in HCS CellMask Blue (Thermo-Fisher, catalog # H32720) or Hoechst 33342 (Invitrogen, catalog # H3570) with Whole Cell Blue (Thermo-Fisher, catalog # 8403502) according to the manufacturers’ protocol. Cells were then washed 3x with PBS, and images were either taken in the DAPI and Cy5 channels using an cellWoRx automated imaging platform (Applied Precision) or in the DAPI and Cy5 channels of an Cellomics ArrayScan VTi (Thermo Fisher).

### Detecting EdU incorporation with Click-iT chemistry

U2OS cells were seeded on sterile coverslips coated with poly-L-lysine (Fisher Scientific, 08-774-383). The doxorubicin treatment protocol described above was followed, with the exception that media was replaced with 10% FBS media 24 and 72 hours after doxorubicin treatment. 10 µM of 5-ethynyl-2’deoxyuridine (EdU, Thermo-Fisher, catalog # A10044) was added to the media 5 days after doxorubicin, and cells were fixed with 4% formaldehyde (Sigma-Aldrich, catalog # F8775) for 15 minutes at room temperature. Cells were then permeabilized with ice-cold methanol for 10 minutes at −20°C. Cells were then washed with PBS 3x, and coverslips were then transferred to a new 24-well plate. Coverslips were stained with HCS CellMask Blue diluted 1:5000 in PBS for 30 minutes, and then washed 3x in PBS. During CellMask incubation, EdU labeling solution was prepared with Click-iT Plus Alexa Fluor 555 Picolyl Azide Toolkit (Thermo-Fisher, catalog # C10642) with the composition according to the manufacturer’s instructions, with 2% CuSO_4_ and 2.5 µM Alexa Fluor 555 picolyl azide. Coverslips were incubated in EdU labeling solution for 30 minutes at room temperature, and then washed 3x in PBS. Coverslips were then mounted onto glass slides using ProLong Gold Antifade Mount. Slides were then imaged on a Nikon spinning-disk confocal microscope.

### Flow cytometry for cell cycle and apoptosis measurements

For all flow cytometry samples, cells were seeded in 15 cm tissue culture dishes 24 hours before doxorubicin treatment. At the indicated time points after doxorubicin treatment, media from the dishes were transferred to 50 mL conical tubes on ice. Cells were then washed with PBS, and afterwards the PBS was added to the respective conical tube (1 tube per plate). Cells were then trypsinized until all cells were detached, and trypsin was subsequently quenched with complete media. Cell suspension was then transferred to the respective conical tube, which was then centrifuged at 1,500 RPM for 5 minutes. The supernatant was then aspirated off, and the cell pellet was resuspended in ice-cold PBS. The cell suspension was then transferred to 1.5 mL Eppendorf tubes, and then centrifuged at 5,000 RPM for 1 minute.

For samples stained for cleaved caspase-3, cleaved PARP, p21, and cyclin B antibody, the pellet supernatant was aspirated, and the cell pellet was resuspended in 4% formaldehyde in PBS and incubated at room temperature for 15 minutes. Formaldehyde solution was removed by centrifuging the cells at 5,000 RPM for 1 minute and removing the supernatant. The cell pellet was then resuspended in ice-cold PBS, and the cell suspension was centrifuged at 5,000 RPM for 1 minute. The supernatant was aspirated off, and cells were resuspended in 90% ice-cold methanol and stored at −20°C until staining. For cleaved caspase-3 and cleaved PARP double staining, cells were washed twice in PBS-T, and then incubated with both antibodies diluted in blocking buffer consisting of 1:1 Odyssey blocking buffer overnight at 4°C, while cells stained for p21 and cyclin B were blocked in blocking buffer for 1 hour at room temperature before proceeding to the overnight primary antibody incubation. For all samples, after the primary antibody step, cells were then washed once with PBS-T, and incubated in respective fluorophore-conjugated secondary antibody diluted in blocking buffer for 1 hour at room temperature for 1 hour. Cells were then washed with PBS-T, and then cleaved caspase-3 and cleaved-PARP stained cells were resuspended in PBS-T for flow cytometry. For the p21 and cyclin B samples, cells were incubated in propidium iodide and RNase A for 1 hour at room temperature before being resuspended in PBS-T for flow cytometry.

For samples stained with BrdU antibody, the pellet supernatant from the ice-cold PBS wash step was aspirated off, and the cell pellet were resuspended in 200 µL ice-cold PBS. 800 µL of ice-cold ethanol was then added drop-by-drop to the Eppendorf tube while vortexing, and samples were then stored at −20°C until staining. Cells were then washed with PBS-T (PBS + 0.5% Tween) and simultaneously permeabilized and denatured with 2 M HCl and 0.5% Triton-X-100 in water for 30 minutes. Cells were washed twice with PBS, and then blocked in blocking buffer consisting of 1:1 Odyssey blocking buffer and PBS-T for 1 hour at room temperature. Blocking buffer was then removed, and cells were then incubated with anti-BrdU antibody diluted in blocking buffer overnight at 4°C. Cells were then washed once in PBS-T, and then were incubated with Alexa Fluor 647 goat anti-mouse antibody diluted in blocking buffer for 1 hour at room temperature. Secondary antibody was removed, and cells were washed once in PBS-T. Cells were then stained with 100 µg/mL propidium iodide with RNase A (Cell Signaling Technology, catalog # 4087) for 1 hour to visualize DNA content.

All flow cytometry measurements were collected on a BD FacsCaliber flow cytometer (BD Biosciences) or a BD LSR II HTS (BD Biosciences).

### Proliferation assay

Cells were seeded at 5,000 cells/well into a tissue culture-treated 96-well plate (Thermo-Fisher, catalog # 165305) 24 hours before doxorubicin treatment. Doxorubicin or DMSO was directly added to culture media, and media was changed to media containing 1% FBS media 4 hours after drug addition. The plate was placed back in the incubator, and then 6 hours after doxorubicin treatment was moved to an incubator connected to an Incucyte microscope (Sartorius). Brightfield images of wells were taken 6, 12, 24, 48, 72, and 96 hours after doxorubicin treatment, and cell number was calculated from these images using ilastik software (Berg et al., 2019). Proliferation values were normalized across all treatments and time points to the 6 hour DMSO cell count values.

### Immunofluorescence measurements for signaling proteins

U2OS cells were seeded into optical bottom 96-well plates (Thermo-Fisher, catalog # 165305) 24 hours before doxorubicin treatment. DMSO or doxorubicin were added directly to the media, and then 4 hours later the media was replaced with DMEM containing 1% FBS, which was maintained for the rest of the experiment. At each indicated timepoint, the media was removed and cells were fixed with 4% formaldehyde in PBS for 15 minutes. Cells were then washed with PBS-Tween (PBS-T) once, and then permeabilized with ice-cold methanol for 10 minutes. Afterwards, cells were again washed once with PBS-T, and then blocked in blocking buffer, which consisted of a 1:1 mix of Odyssey blocking buffer (Licor) and PBS-T. Cells were blocked for 1 hour at room temperature.

Afterwards, the blocking buffer was removed and replaced with primary antibody solution consisting of antibody diluted in blocking buffer. Cells were incubated in primary antibody solution overnight at 4°C on a shaker. The primary antibody solution was then removed, and cells were washed once with PBS-T. Cells were then incubated with a secondary antibody solution containing both goat anti-mouse conjugated to Alexa Fluor 488 and goat anti-rabbit conjugated to Alexa Fluor 647 diluted in blocking buffer. Cells were incubated in secondary antibody solution overnight at 4°C on a shaker, and then washed once with PBS-T. To stain both the nuclear and the cytosolic compartments, cells were incubated with either 1:1000 Whole Cell Blue with 1:10,000 Hoechst 33342 in PBS or 1:5000 HCS CellMask Blue in PBS for 1 hour at 4°C. Cells were then washed with PBS, and wells were replaced with PBS for imaging. Cells were imaged in the DAPI, GFP, and Cy5 channels on either a cellWoRx automated imaging platform (Applied Precision) or in the DAPI and Cy5 channels of a Cellomics ArrayScan VTi (Thermo Fisher).

### Signal and response data processing

To generate the signaling dataset, raw immunofluorescence images were processed in Fiji (Schindelin et al., 2012) and subsequently run through a pipeline in CellProfiler for intensity quantification (version 4.0.6) (Stirling et al., 2021). Mean intensity of the nuclear and cytoplasmic compartments (where applicable) of total and phospho-proteins at all time points were normalized to the 2 µM doxorubicin value at the 6 hour time point across a single experiment. The mean of all biological replicates across experiments for each signal was used for subsequent data visualization and modeling.

The response dataset consisted of normalized fold-change data for the proliferation measurements, and percent positive as gated in flow cytometry for the rest of the response measurements collected. The mean was calculated for each individual response and time point across all biological replicates and experiments. Then, percent positive measurements were divided by 100 to scale values between 0 and 1, and a logit transformation was applied to all data points. Values of 0 were converted to 0.00001 for the logit transformation.

For principal component analysis (PCA), the normalized signals data was shaped into a 4×162 matrix, with rows representing the 4 distinct treatments and columns representing individual signals (27) and timepoints (6). The z-score was taken of this 4×162 matrix, which was used for principal component analysis.

For tensor PLSR, the raw signal data (X) was shaped into a 4×6×27 tensor, with mode 1 representing treatments (n = 4), mode 2 representing time points (n = 6), and mode 3 representing each individual signal (n = 27). The response data (Y) was shaped into a 4×6×10 tensor, with the same modes 1 and 2, and mode 3 representing individual responses (n = 10).

### Tensor PLSR and VIP score calculation

The X and Y tensors were mean-centered across mode 1 and variance scaled across modes 2 and 3. The tensor PLSR model was constructed in MATLAB 2019b using the N-way Toolbox (version 1.8.0.0) as described in (Chitforoushzadeh et al., 2016). All scripts and packages are available for download in supplementary material (S#).

The VIP scores were calculated as described in (Favilla et al., 2013, 2014) using the Multi-Way VIP package in MATLAB (Favilla et al., 2013).

### Principal component analysis

The z-scored signaling matrix of 4 x 162 was run through the pca() function in MATLAB using three principal components. Scores and loadings were plotted on principal components one and two, as these explained the majority of the variance in the signals (> 90%). All of the scores and loadings of PC 2 were multiplied by −1 to improve the interpretability in comparison to the t-PLSR results. See supplemental code (S#) for more details.

### Inhibitor co-treatment experiments

Cells were seeded at 5,000 cells/well into an optical bottom 96-well plate (Thermo-Fisher, catalog # 165305) 24 hours before doxorubicin treatment. For the 0-12 hour inhibitor pulse, cells were treated with either vehicle (DMSO) or doxorubicin in addition to 10 µM SP600125, 10 µM PD98059, 10 µM SB203580, caffeine (5 mM for U2OS cells and 1 mM for OVCAR-8 cells), or vehicle (DMSO). Cells were maintained in this media for 4 hours, and then media was replaced with 1% FBS media containing either the respective inhibitor or vehicle, media was replaced with 10% FBS media 1 and 3 days after doxorubicin treatment, and inhibitor was maintained for the entire experiment for the results seen in figure 5E-G. For figures 6 and S10, 12 hours after the addition of doxorubicin, the media was replaced with 1% FBS media containing no inhibitors or vehicle. For the 12-24 hour inhibitor pulse, cells were treated with either vehicle (DMSO) or doxorubicin for 4 hours in the absence of inhibitors, and then the media was replaced with 1% FBS media. 12 hours after the addition of doxorubicin, the media was replaced with media containing the above inhibitors at the listed concentrations. Media was replaced with 10% FBS media 1 day and 3 days after the addition of doxorubicin for both the 0-12 and the 12-24 hour inhibitor condition. 5 days after the addition of doxorubicin, 10 µM BrdU was added to all wells, and 24 hours later plates were harvested for BrdU immunofluorescence, which was performed as described above.

### ELISA for IL-6 and IL-8

U2OS cells were seeded at 5,000 cells/well into a 96-well plate 24 hours before doxorubicin treatment. Cells were treated with either vehicle or doxorubicin for 4 hours, and then media was replaced with drug-free, 1% FBS media. 24 hours after the addition of doxorubicin, media was replaced with 10% FBS media. Three days after doxorubicin addition, 10% FBS media containing either vehicle (DMSO), 10 µM SP600125, or 10 µM PD98059 was used to replace the growth media. Five days after doxorubicin treatment, the media was changed in wells to media without serum, and six days after doxorubicin treatment, media was transferred to a v-bottom 96-well plate, and stored at −80°C. ELISA assays for IL-6 (Invitrogen, catalog # KHC0061) and IL-8 (Invitrogen, catalog # KHC0081) levels in the media samples were conducted according to the manufacturer’s protocol.

### Immunoblotting

Cells were treated as indicated in 6-well plates. At the given time points, the culture media was aspirated and cells were immediately lysed on ice. For the siRNA knockdown blots, RIPA buffer [50 mM Tris-HCl (pH 8), 150 mM NaCl, 0.1% sodium dodecyl sulfate (SDS), 0.5% sodium deoxycholate, 1% Triton-X] was used, while for the rest of the shown blots high SDS (0.7%) lysis buffer [10 mM Tris (pH 7.5), 50 mM NaCl, 0.5% NP-40, 0.5% sodium deoxycholate, 0.7% SDS, 10 mM iodoacetamide] was used. Both types of lysis buffer were supplemented with phosSTOP (Roche, catalog # 4906837001) and cOmplete (Roche, catalog # 5892791001) tabs. 100 µL of lysis buffer was added to each well after aspiration, and afterwards was evenly distributed within the well. The plate was then incubated on ice for 5 minutes, and a cell scraper was used to scrape remaining cells while the plate was still on ice. The lysates were transferred to a cold Eppendorf tube and subsequently sonicated and centrifuged at 18,000 x g for 10 minutes at 4°C for the RIPA buffer lysates. For the high SDS lysates, lysates were vortexed for 15 seconds and then put back on ice for 10 minutes after the sonication step. This vortexing step was repeated twice for a total of 30 minutes, and then lysates were centrifuged at 18,000 x g for 10 minutes at 4°C. After centrifugation, the supernatants were transferred over to a new, pre-chilled Eppendorf tube and stored at −20°C.

The protein concentration of lysates was measured using a BCA assay (Thermo-Fisher, catalog # 23225), and the protein concentration was normalized across all samples. Samples were boiled for 5 minutes in 1x sample buffer [34 mM Tris, pH 6.8, 7% glycerol, 500 mM 2-mercaptoethanol, 1.6% SDS], and then loaded on a 10% SDS PAGE gel. After the samples ran through the gel at 100 V, a wet transfer step was run at 100 V for 1 hour at 4°C to transfer protein to a 0.22 µM nitrocellulose membrane. The quality of the transfer was checked with a Ponceau staining (Sigma-Aldrich, catalog # P7170), and after destaining the membranes were blocked in a blocking buffer consisting of a 1:1 mix of Intercept Blocking Buffer (Licor, catalog # 927-70001) and PBS-T (PBS + 0.05% Tween) at room temperature for 1 hour on a shaker. Primary antibody was diluted in blocking buffer, and incubated with the membrane overnight at 4°C on a shaker. The membrane was then washed five times quickly with PBS-T, and then washed three times for 5 minutes each in PBS-T on a shaker at room temperature. Membranes were then incubated with secondary antibody (Licor) diluted in blocking buffer for 1 hour at room temperature on a shaker, and afterwards the same wash steps as described above were performed, with the exception that the last wash step was conducted with PBS instead of PBS-T. Membranes were then kept in PBS, and then imaged in the 680 and 800 channel of an Odyssey CLx machine (Licor). The mean, background subtracted signal intensity of respective bands were quantified in Image Studio Lite (Licor, version 5.2), and intensity values were used for described ratio calculations.

For blots of phospho and total c-Jun, JNK, the blot was probed first using the phospho antibody, and then the membrane was stripped with 1x NewBlot nitrocellulose stripping buffer (Licor, catalog # 928-40030), washed 5x quickly in PBS, and then washed three times for 10 minutes each in PBS on a shaker at room temperature. The membrane was then blocked again in 1:1 Odyssey blocking buffer to PBS-T, and reprobed with the total antibody on a shaker overnight at 4°C. The total MK2 blots were stripped and reprobed for p38 as described above on the same membrane, which was also done for the p-MK2 blots with p-p38.

### siRNA transfection

Non-targeting (catalog #AM4636) was purchased from Invitrogen, and Jun #1 (catalog # s7658), and Jun #2 (catalog # s7659) siRNAs were purchased from Life Technologies. U2OS cells were plated in 6-well plates at a density of 150,000 cells/well 24 hours prior to transfection. Lipofectamine RNAiMAX was purchased from ThermoFisher Scientific (catalog # 13778150), and used according to the manufacturer’s protocol to dose cells with 10 nM of siRNA-Lipofectamine duplex. Cells were incubated in siRNA/Lipofectamine mixture for 24 hours, and then each well was seeded into a 96-well plate at a density of 5,000 cells/well in triplicate. A concurrent 6-well plate transfected with siRNAs had media changed 24 hours post-transfection, and cells were harvested for immunoblotting 48 hours after transfection.

Cells in the 96-well plate were treated with doxorubicin 24 hours after seeding, and media was changed to media containing 1% FBS 4 hours after doxorubicin treatment. 24 hours after doxorubicin treatment, media was changed to complete media containing 10% FBS. Immunofluorescence for BrdU was conducted six days after doxorubicin treatment, as described above.

### Cloning and Tol2-mediated stable transfection

pAK-Tol2-TRE-JunDN-HA-T2A-GFP (pTRE-JunDN-HA-T2A-GFP) construct was assembled using Gibson assembly with NEBuilder HiFi DNA Assembly Master Mix (NEB, catalog # E2621) and inserted into a lentiviral pLVX-CMV empty backbone. pMIEG3-JunDN was a gift from Alexander Dent (Addgene plasmid # 40350; http://n2t.net/addgene:40350; RRID:Addgene_40350) (Z.-Y. Wang et al., 2005), and was used as a template to amplify JunDN-HA and GFP fragments used in the assembly. JunDN-HA-T2A-GFP was amplified from the pLVX-CMV backbone using Phusion Mastermix (NEB, catalog # M0531), and inserted into the empty backbone of pAK_Tol2_TRE_Puro (Addgene, plasmid #130259) with digestion and ligation cloning into the NheI (NEB, catalog # R3131) and EcoRI (NEB, catalog # R3101) sites. GFP amplified from pLVX-CMV-JunDN-HA-T2A-GFP was inserted as described above into the same backbone for the TRE-GFP control construct. Positive colonies for both plasmids were confirmed with Sanger sequencing.

U2OS cells were seeded at a density of 200,000 cells/well into a 6-well plate 24 hours before transfection. Cells were co-transfected with 1.25 µg pCMV-Tol2 and 1.25 µg of either pAK-Tol2-TRE-JunDN-HA-T2A-GFP or pAK-Tol2-TRE-GFP (pTRE-GFP) with Lipofectamine 2000 (ThermoFisher Scientific, catalog # 11668019) according to the manufacturer’s instructions at a 1:2 DNA (µg) to Lipofectamine 2000 reagent (µL) ratio. Media was replaced 24 hours later, and cells were selected with 2 µg/mL puromycin (Invivogen, catalog # ant-pr-1) 96 hours post-transfection for 3 days. Cells were then induced with 1 µg/mL doxycycline for 3 days, and GFP positive cells were harvested by FACS using a BD FACSAria III machine (BD Biosciences) for further experiments.

### c-Jun dominant negative experiments

U2OS cells stably expressing either the pTRE-GFP or the pTRE-JunDN-HA-T2A-GFP construct and were FACS-sorted for GFP positivity as described above were seeded in a 96-well plate at 1,000 cells/well 96 hours before doxorubicin treatment. 72 hours before doxorubicin treatment, media was changed to complete media with or without 1 µg/mL doxycycline. 24 hours before doxorubicin treatment, media was changed again to complete media with or without 1 µg/mL doxycycline. Doxorubicin or DMSO was added directly to the wells, and media was changed to media containing 1% FBS with or without 1 µg/mL doxycycline. 12 hours after doxorubicin treatment, media was changed to media containing 1% FBS without doxycycline, and 24 and 72 hours after doxorubicin treatment, the media was changed to complete media containing 10% FBS. Immunofluorescence for BrdU was conducted as described above.

### Retrovirus production and transduction

pSIRV-AP-1-mCherry was a gift from Peter Steinberger (Addgene plasmid # 118095; http://n2t.net/addgene:118095; RRID:Addgene_118095) (Jutz et al., 2016), and pUMVC was a gift from Bob Weinberg (Addgene plasmid # 8449; http://n2t.net/addgene:8449; RRID:Addgene_8449) (S. A. Stewart et al., 2003). HEK293T cells were co-transfected with 15 µg pSIRV-AP-1-mCherry, 10 µg pUMVC, and 2.5 µg pCMV-VSVG using the calcium phosphate transfection system (Clonetech), and media was changed 24 hours later. Media was harvested 24 hours after the media changed, and filtered through a 0.45 µm filter with a syringe. Filtered media was given to U2OS cells for 24 hours, and two weeks post-transduction, mCherry positive cells were harvested by FACS using a BD FACSAria III machine (BD Biosciences) for further experiments.

To generate U2OS cells containing both AP-1-mCherry and JunDN-IRES-eGFP, FACS-sorted AP-1-mCherry cells from above were infected with pMIEG3-JunDN construct packaged in retrovirus. pMIEG3-JunDN was packaged with HEK293Ts as described above, and filtered media containing virus was diluted 1:1 into complete media. U2OS cells were incubated with virus for 24 hours, and two weeks later, a GFP high population was harvested by FACS for further experiments using a BD FACSAria III machine (BD Biosciences).

### AP-1-mCherry experiments

U2OS cells stably transfected with the AP-1 mCherry reporter were seeded into 6-well plates at a density of 150,000 cells/mL 24 hours before treatment. Cells were treated with either DMSO or doxorubicin for 4 hours, and then media was replaced with DMEM containing 1% FBS. For the entire experiment, cells were co-treated with either DMSO, 10 µM SP600125, or 10 µM PD98059. AP-1-mCherry cells co-expressing JunDN-IRES-GFP were also treated with either DMSO or doxorubicin. As a positive control, AP-1-mCherry cells were treated with 100 nM phorbol 12-myristate 13-actetate (PMA, Sigma-Aldrich, catalog # 79346) for 24 hours and was administered at the same time as the doxorubicin. Cells were harvested 24 hours after treatment, and flow cytometry was performed using a BD LSR II HTS flow cytometer (BD Biosciences).

### OVCAR-8 viability curve

OVCAR-8 cells were seeded at a density of 5,000 cells/well into a 96 well plate 24 hours before treatment. Doxorubicin was added to wells in triplicate for a final concentration of: 0.795 nM, 15.6 nM, 31.25 nM, 62.5 nM, 125 nM, 250 nM, 500 nM, 1000 nM, and 2000 nM. Cells were then incubated in DMSO vehicle or these doses of doxorubicin for 4 hours, and then media was changed to complete media containing 10% FBS. Media was changed again 24 hours after the start of doxorubicin treatment, and 96 hours after the start of doxorubicin treatment, ATP levels were measured with the CellTiter-Glo Luminescent Cell Viability assay (Promega, catalog # G7570) according to the manufacturer’s protocol, and luminescence values were normalized to the vehicle control.

### Software

For image processing, either Fiji or NIS Viewer were used. CellProfiler was used for downstream image quantification, and FlowJo v10 was used to process flow cytometry-based response data. For Incucyte-based proliferation data, ilastik was used to count the number of cells in brightfield images. MATLAB 2019b was used for modeling and PCA analysis, and Python 3.7 was used to analyze and compile CellProfiler outputs as well as generate boxplots. Statistics and bar plots were computed and made using Graphpad Prism 8.4.1. All cartoon illustrations were made directly in Adobe Illustrator, or by using BioRender under an academic license. Publication licenses are available upon request.

---

**Figure S1:**
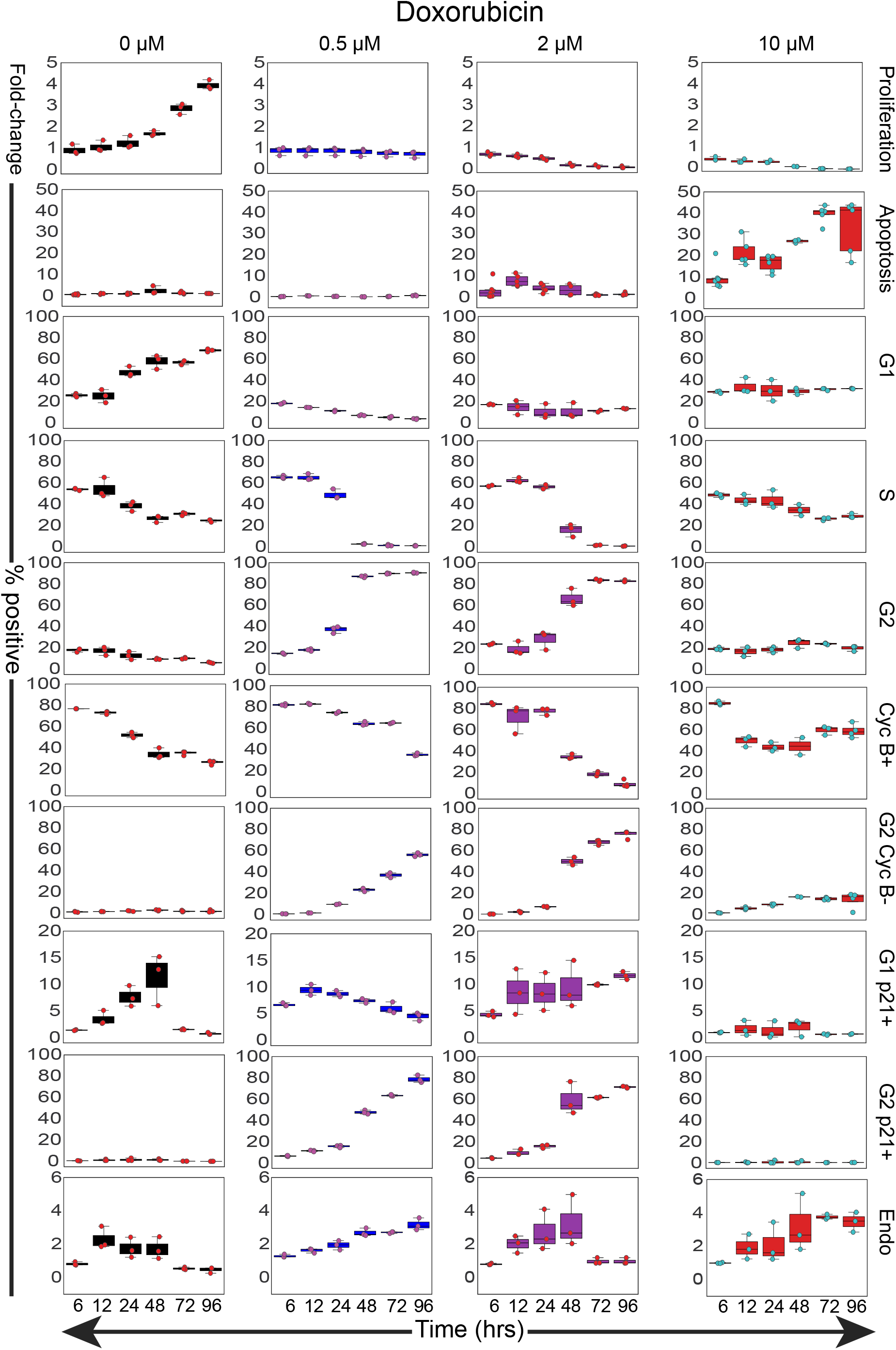
Boxplots of raw response data. Boxplots of raw response data vs. time. Y-axis values represent fold-change for proliferation values, and percent positive values for responses measured by flow cytometry. Overlaid dots represent individual biological replicates.

**Figure S2:**
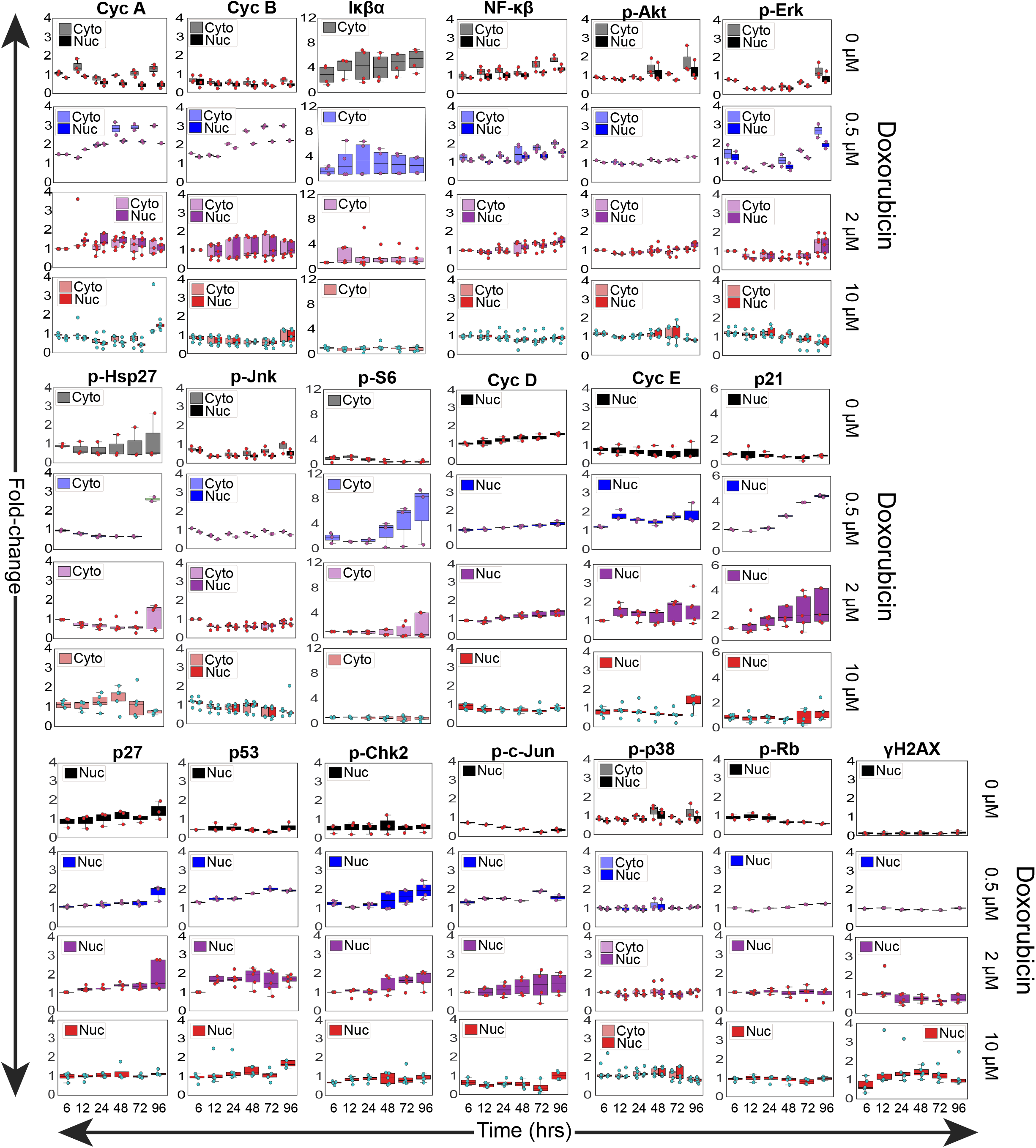
Boxplots of raw signals data. Boxplots of raw signals data vs. time. Y-axis values represent fold-change of the mean intensity normalized to the 2 µM dose at the 6 hour timepoint for both the nuclear (Nuc) and cytoplasmic (Cyto) compartments. Overlaid dots represent individual biological replicates.

**Figure S3:**
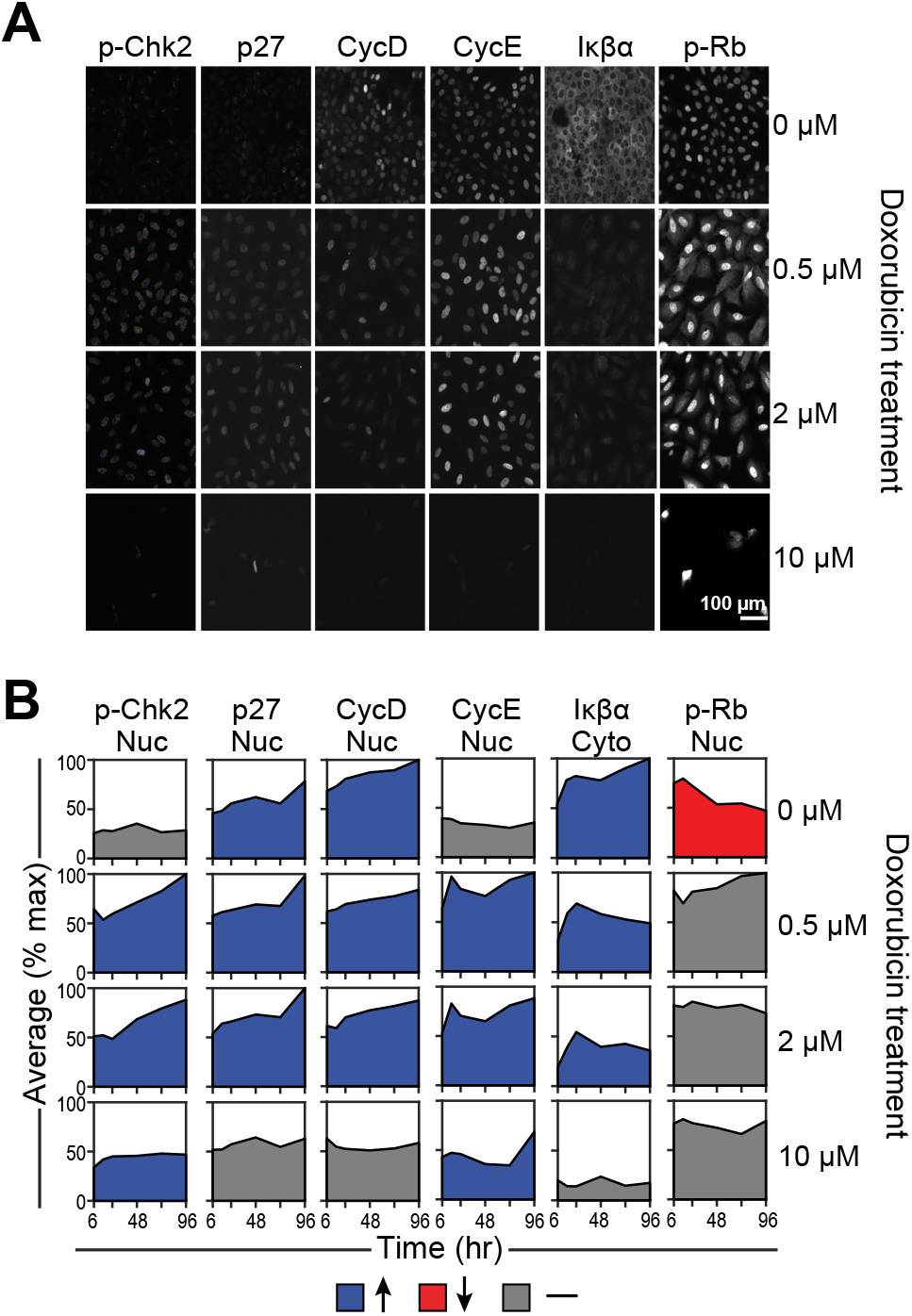
Quantification of remaining signaling measurements. **A)** Representative immunofluorescence images of remaining signals not shown in figure 3. **B)** Quantification of the mean fluorescence intensity (normalized to the maximum value across time and drug treatments) over time in either the nuclear or cytoplasmic compartment, depending on the given protein measured. Tick marks on the x-axis represent 6, 24, 48, 72, and 96 hrs.

**Figure S4:**
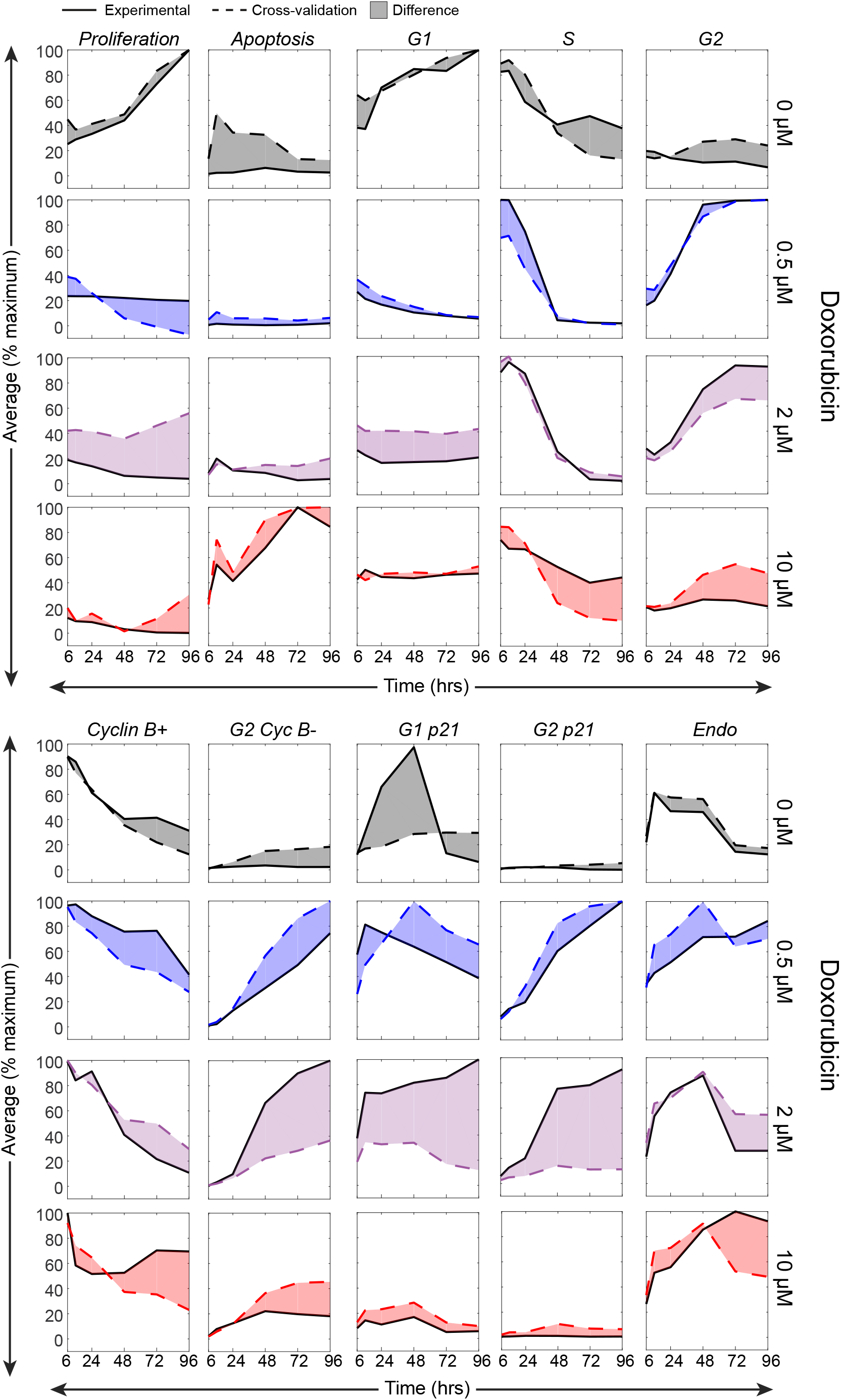
The 2 µM doxorubicin dose cross-validation values diverge the most from experimental values. The mean fluorescence intensity (normalized to the maximum value across time and drug treatments) vs. time is plotted for each of the predicted responses, with the solid lines representing the experimental values, the dotted lines representing the cross-validation predictions, and the shaded area highlighting the difference between the two curves.

**Figure S5:**
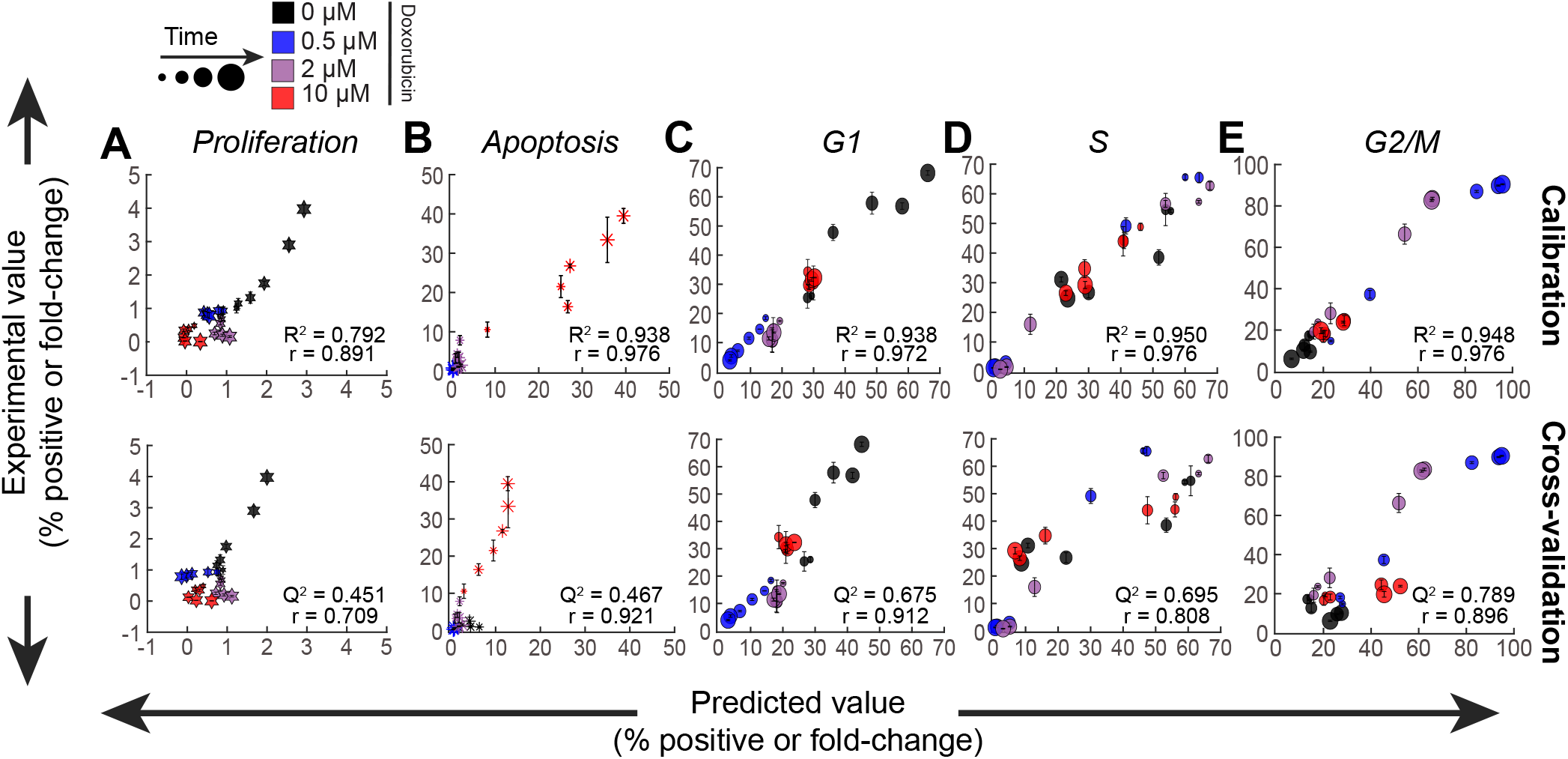
Experimental vs. predicted values on an individual response basis. Scatterplots of experimental vs. predicted values for **A)** Proliferation **B)** Apoptosis **C)** G1 **D)** S and **E)** G2/M responses from the calibration and the cross-validation model. R^2^, Q^2^, and Pearson correlation values are also shown.

**Figure S6:**
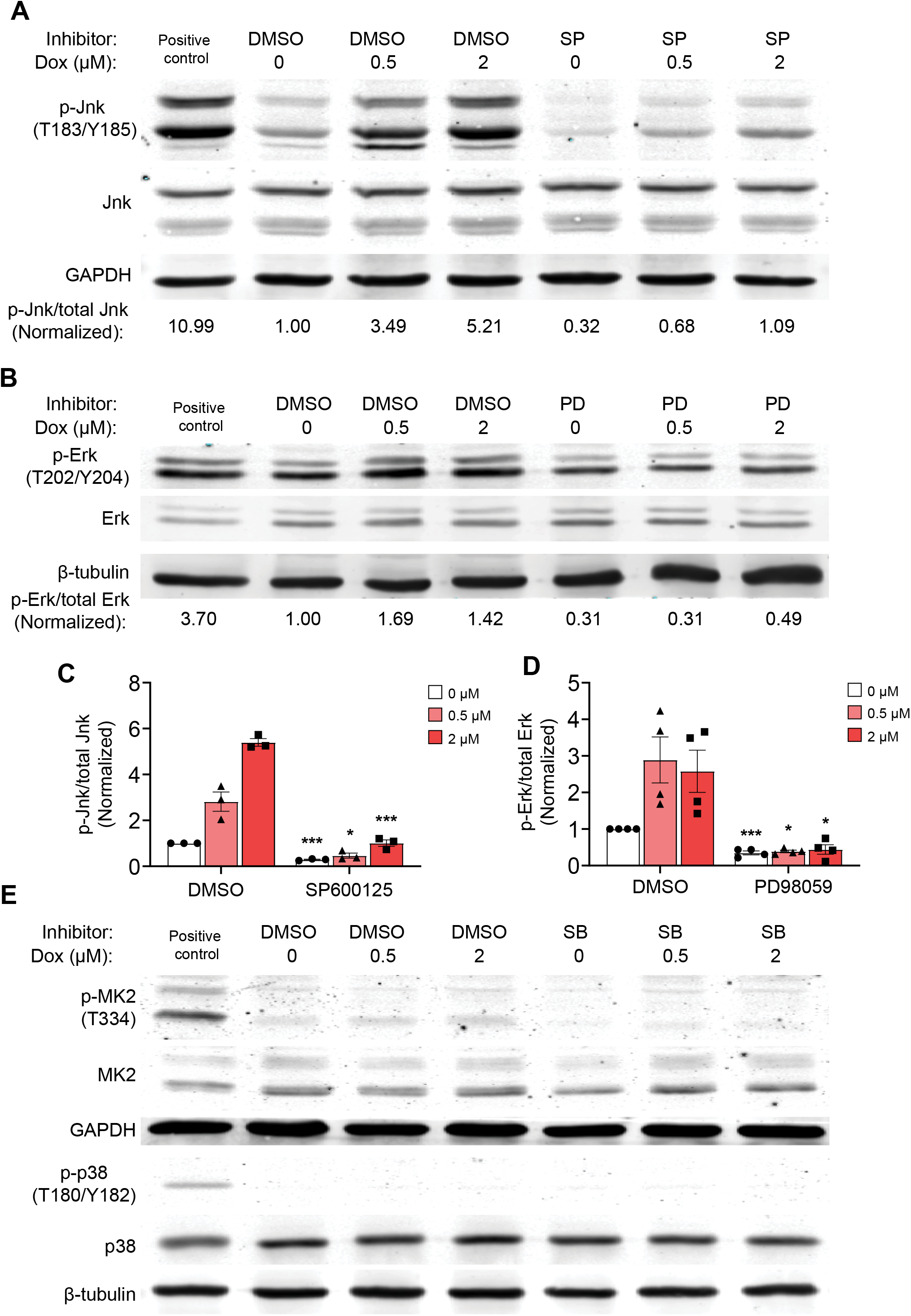
Western blotting confirms that inhibitors decrease phosphorylation of expected downstream targets. **A)** Representative western blot of p-JNK (T183/Y185), JNK, and GAPDH in U2OS cells that were treated with either vehicle (0 µM) or doxorubicin (0.5 µM and 2 µM) for four hours, and cells were either co-treated with vehicle (DMSO), SP600125 (SP, JNKi) or PD98059 (PD, Meki. Samples were run with a positive control of U2OS cells treated with 25 µg/mL anisomycin for 15 minutes. **B)** Representative western blot of p-Erk (T202/Y204), Erk1/2, and β-tubulin in U2OS samples in the same treatment conditions as subpanel A. Samples were run with a positive control of 100 nM PMA treatment for 30 minutes. **C)** Quantification of p-JNK to total JNK ratio, normalized to DMSO, 0 µM condition. Bars represent mean ± SEM of three biological replicates. **D)** Quantification of p-Erk to total Erk1/2 ratio, normalized to DMSO, 0 µM condition. Bars represent mean ± SEM of four biological replicates. **E)** Representative western blots of p-MK2 (T334), MK2, p-p38 (T180/Y182), p38, GAPDH, and β-tubulin in in U2OS cells that were treated with either vehicle (0 µM) or doxorubicin (0.5 µM and 2 µM) for four hours, and cells were either co-treated with vehicle (DMSO), or SB203580 (SB, p38i). Samples were run with a positive control of U2OS cells treated with 25 µg/mL anisomycin for 15 minutes. Western blots are representative of two biological replicates. ***: p < 0.001 *: p < 0.05 with a two-tailed t-test with Bonferroni correction tested vs. the respective doxorubicin dose in the vehicle inhibitor control in subpanels C and D.

**Figure S7:**
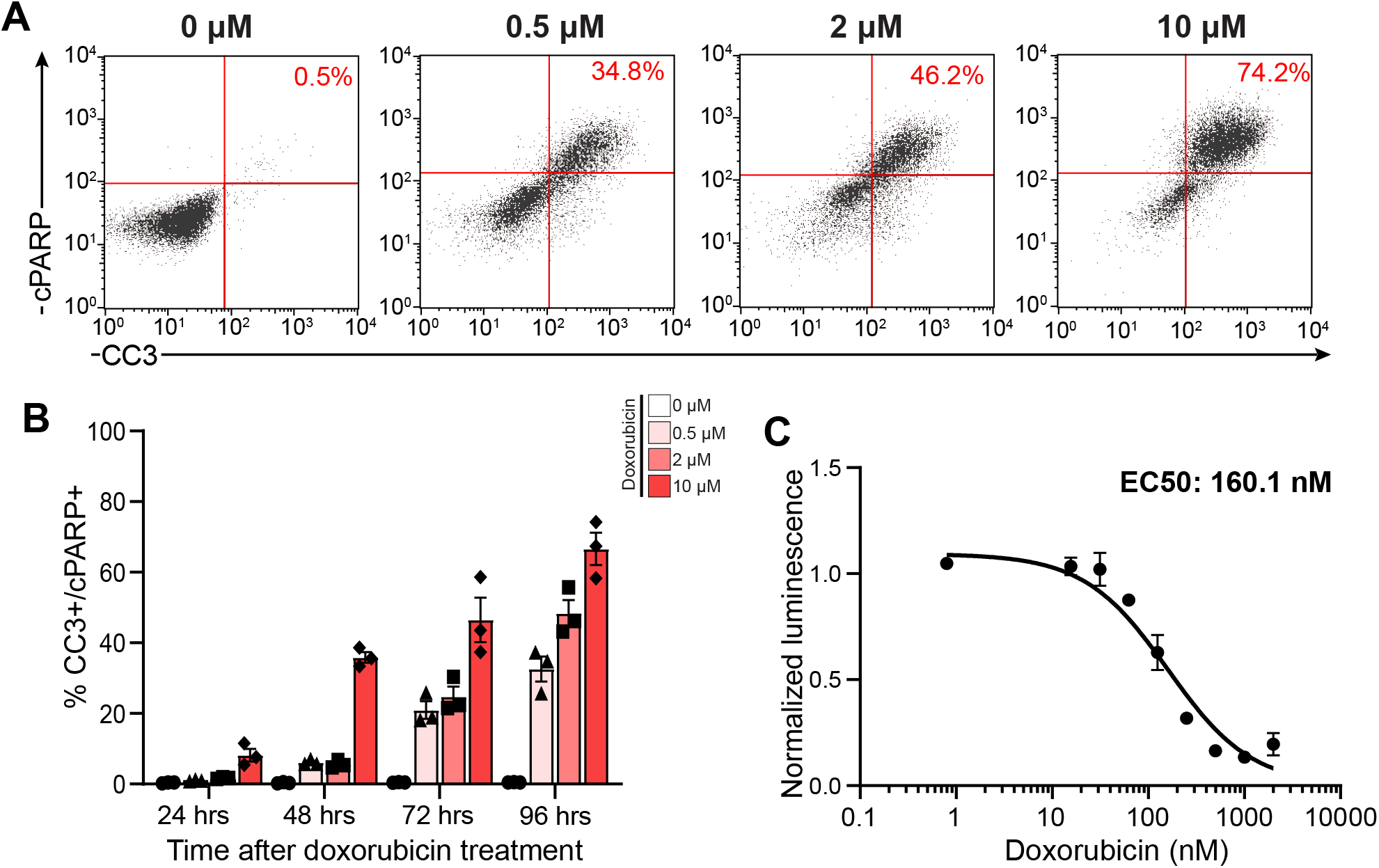
OVCAR-8 cells undergo dose-dependent apoptosis after doxorubicin treatment. **A)** Representative flow cytometry plots of cleaved PARP (cPARP) vs. cleaved caspase-3 (CC3) 96 hours after doxorubicin treatment. **B)** Quantification of CC3/cPARP double positivity over time at varying doses of doxorubicin. Bars represent the mean ± SEM of three biological replicates. **C)** Viability curve of OVCAR-8 cells using total ATP measurements 96 hours after doxorubicin treatment. All points were normalized to no treatment control, each point represents the mean ± SEM of three biological replicates.

**Figure S8:**
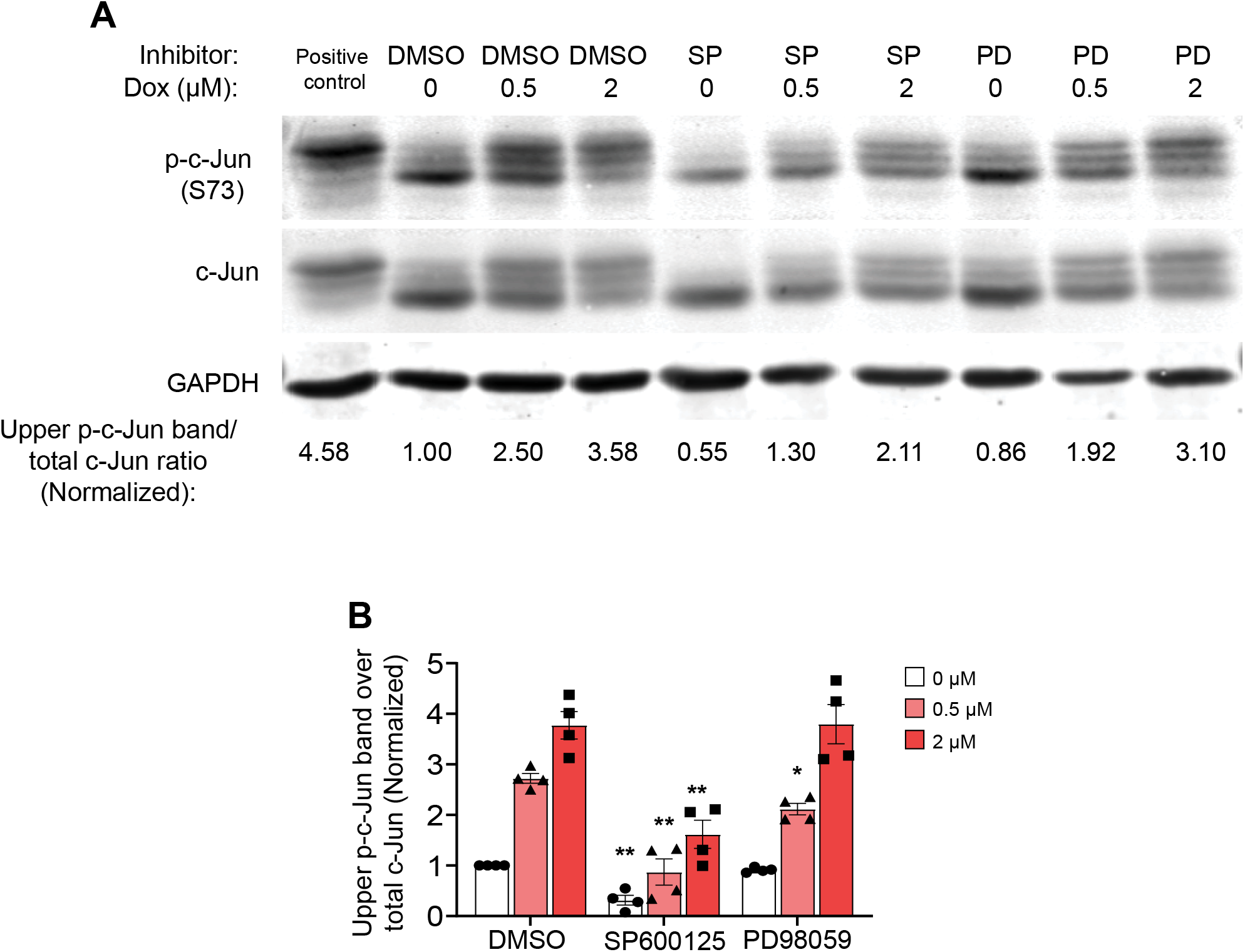
Immunoblotting reveals a decrease in p-c-Jun(S73) at certain doses of doxorubicin with a JNK or Mek inhibitor co-treatment. **A)** Representative western blot probed for p-c-Jun (S73), c-Jun, and GAPDH in U2OS cells that were treated with doxorubicin for four hours, and cells were either co-treated with vehicle (DMSO), or with either 10 µM SP600125 (SP, JNKi) or 10 µM PD98059 (PD, Meki). Cells were lysed 6 hours after doxorubicin treatment for western blotting. Samples were run with a positive control of U2OS cells treated with 25 µg/mL anisomycin for 15 minutes. **B)** Quantification of the ratio of the upper p-c-Jun(S73) band to total c-Jun. Bars represent the mean ± SEM of four biological replicates, and bars represent SEM. **: p < 0.01 *: p < 0.05 with a two-tailed t-test with Bonferroni correction tested vs. the respective doxorubicin dose in the vehicle inhibitor control.

**Figure S9:**
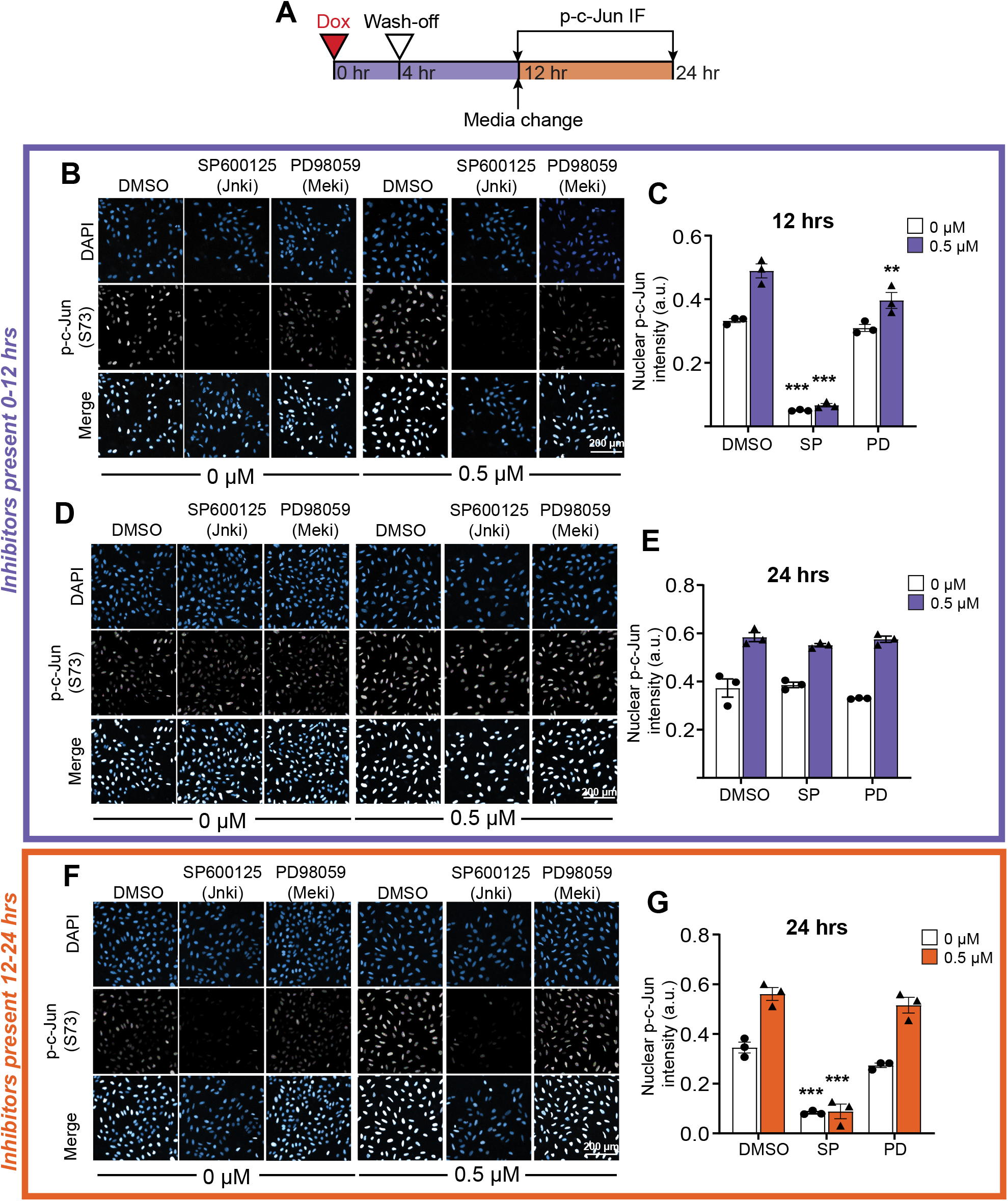
Time course of p-c-Jun(S73) show inhibitor-dependent decreases in phosphorylation levels in the first 12 hours after doxorubicin treatment in U2OS cells. **A)** Schematic of the experiment performed. **B)** Representative images of cells immuno-stained for p-c-Jun(S73) and DAPI co-treated with inhibitors the first 12 hours of the experiment, and fixed at the 12 hour timepoint. **C)** Quantification of the nuclear mean p-c-Jun73 intensity 12 hour timepoint. **D)** Representative images of cells immune-stained for p-c-Jun(S73) and DAPI co-treated with inhibitors the first 12 hours of the experiment, and fixed at the 24 hour timepoint. **E)** Quantification of the nuclear mean p-c-Jun73 intensity for the 24 hour timepoint after early inhibitor. **F)** Representative images of cells immune-stained for p-c-Jun(S73) and DAPI co-treated with inhibitors the 12-24 hour window of the experiment, and fixed at the 24 hour timepoint. **G)** Quantification of the nuclear mean p-c-Jun73 intensity for the 24 hour timepoint after late inhibitor. For subpanels C, E, and G, bars represent the mean ± SEM of three biological replicates. ***: p < 0.001 **: p < 0.01 with a two-way ANOVA and post-hoc Dunnett’s test when compared to the DMSO vehicle control at the same doxorubicin dose.

