## Supplemental information for "Biphasic JNK–Erk Signaling Separates Induction and Maintenance of Cell Senescence after DNA Damage"

### Netterfield et al., Figure S1

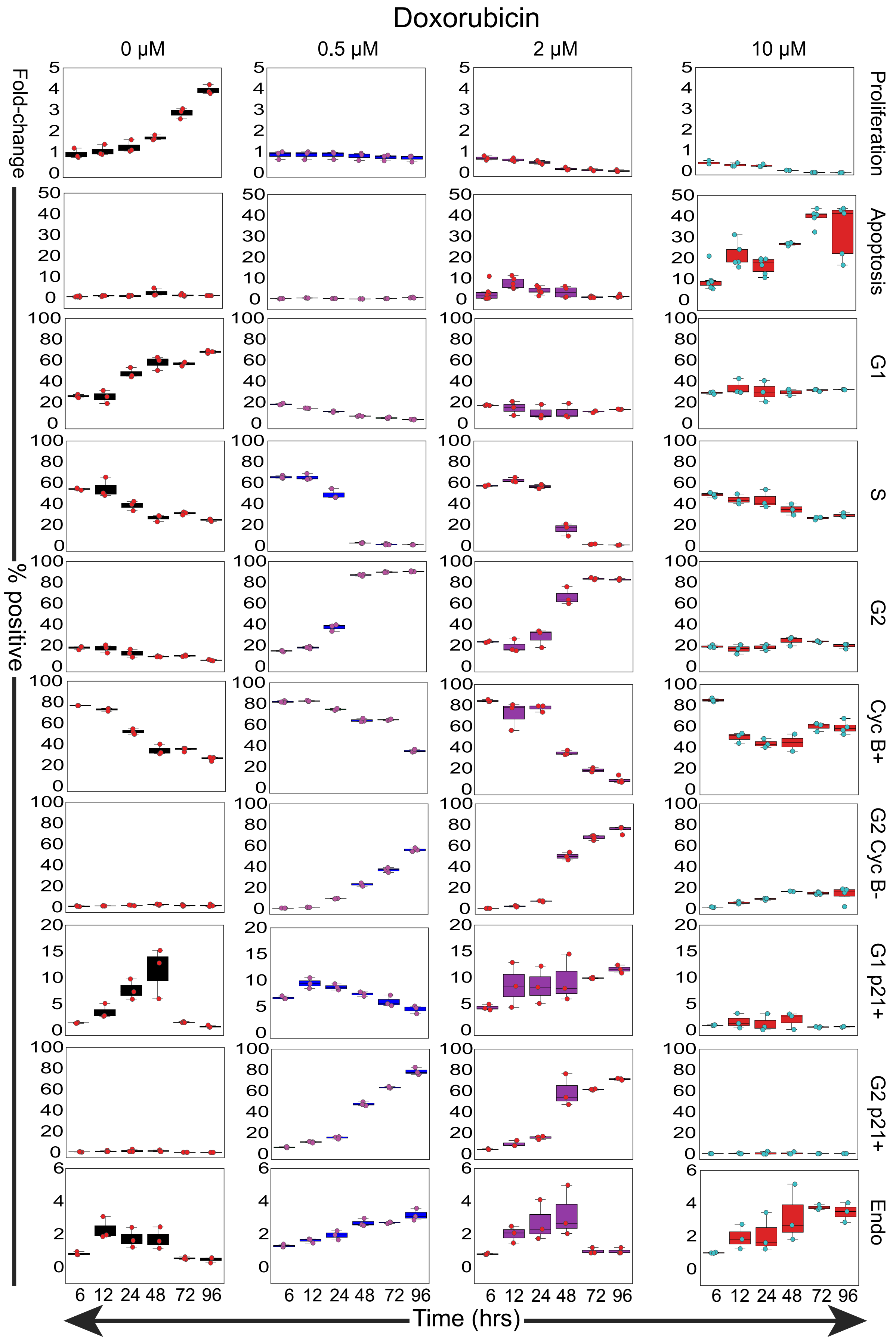

**Figure S1: Boxplots of raw response data.**

Boxplots of raw response data vs. time. Y-axis values represent fold-change for proliferation values, and percent positive values for responses measured by flow cytometry. Overlaid dots represent individual biological replicates.

**Netterfield et al., Figure S2**

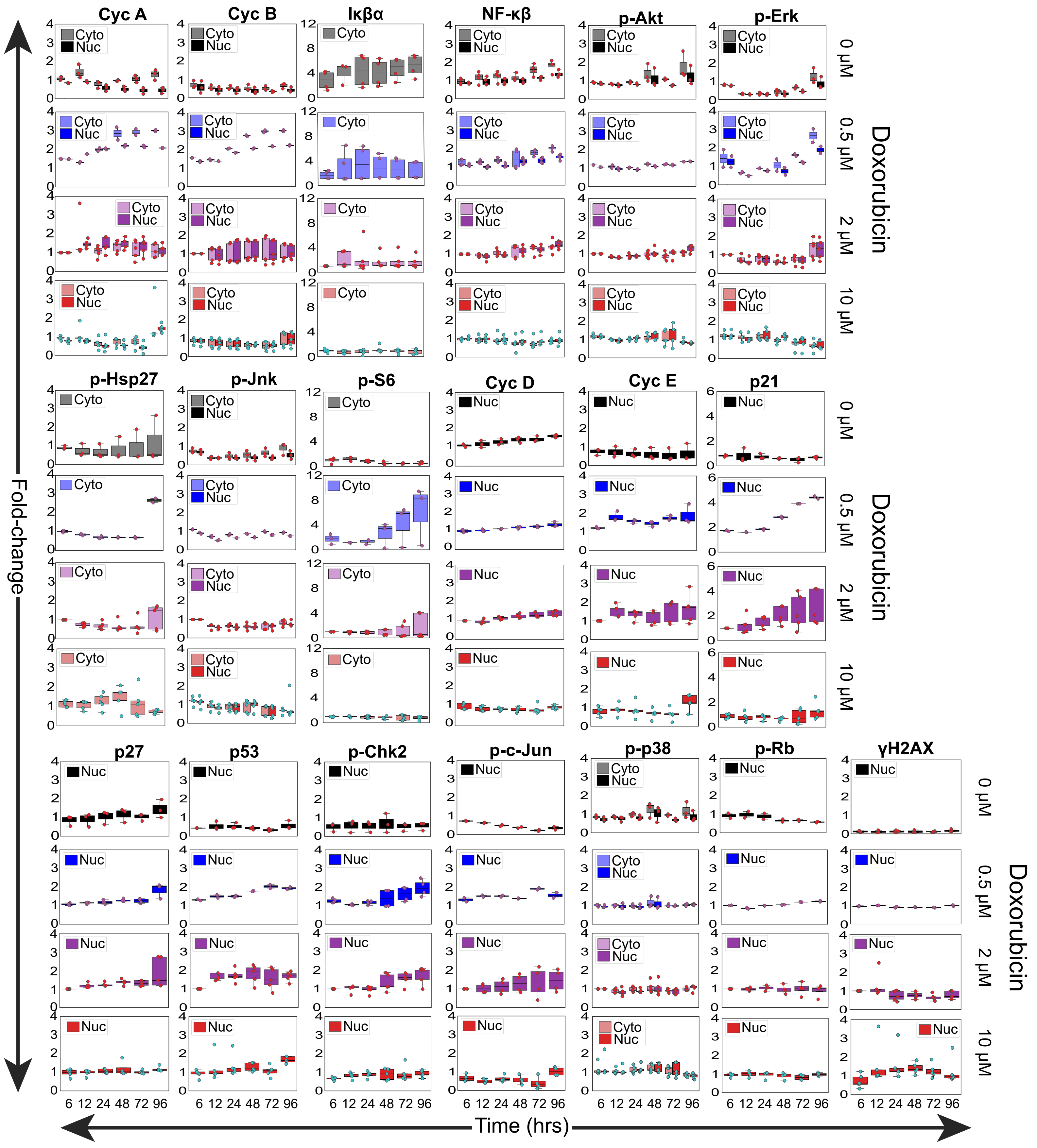

**Figure S2: Boxplots of raw signals data.**

Boxplots of raw signals data vs. time. Y-axis values represent fold-change of the mean intensity normalized to the 2 μM dose at the 6 hour timepoint for both the nuclear (Nuc) and cytoplasmic (Cyto) compartments. Overlaid dots represent individual biological replicates.

### Netterfield et al., Figure S3

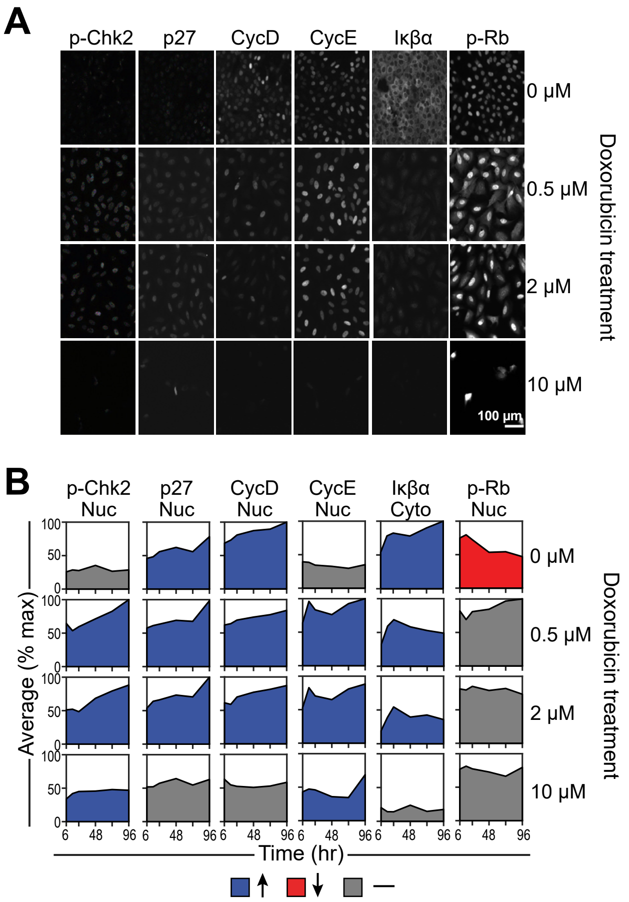

**Figure S3: Quantification of remaining signaling measurements.**

**A)** Representative immunofluorescence images of remaining signals not shown in figure 3.
**B)** Quantification of the mean fluorescence intensity (normalized to the maximum value across time and drug treatments) over time in either the nuclear or cytoplasmic compartment, depending on the given protein measured. Tick marks on the x-axis represent 6, 24, 48, 72, and 96 hrs.

### Netterfield et al., Figure S4

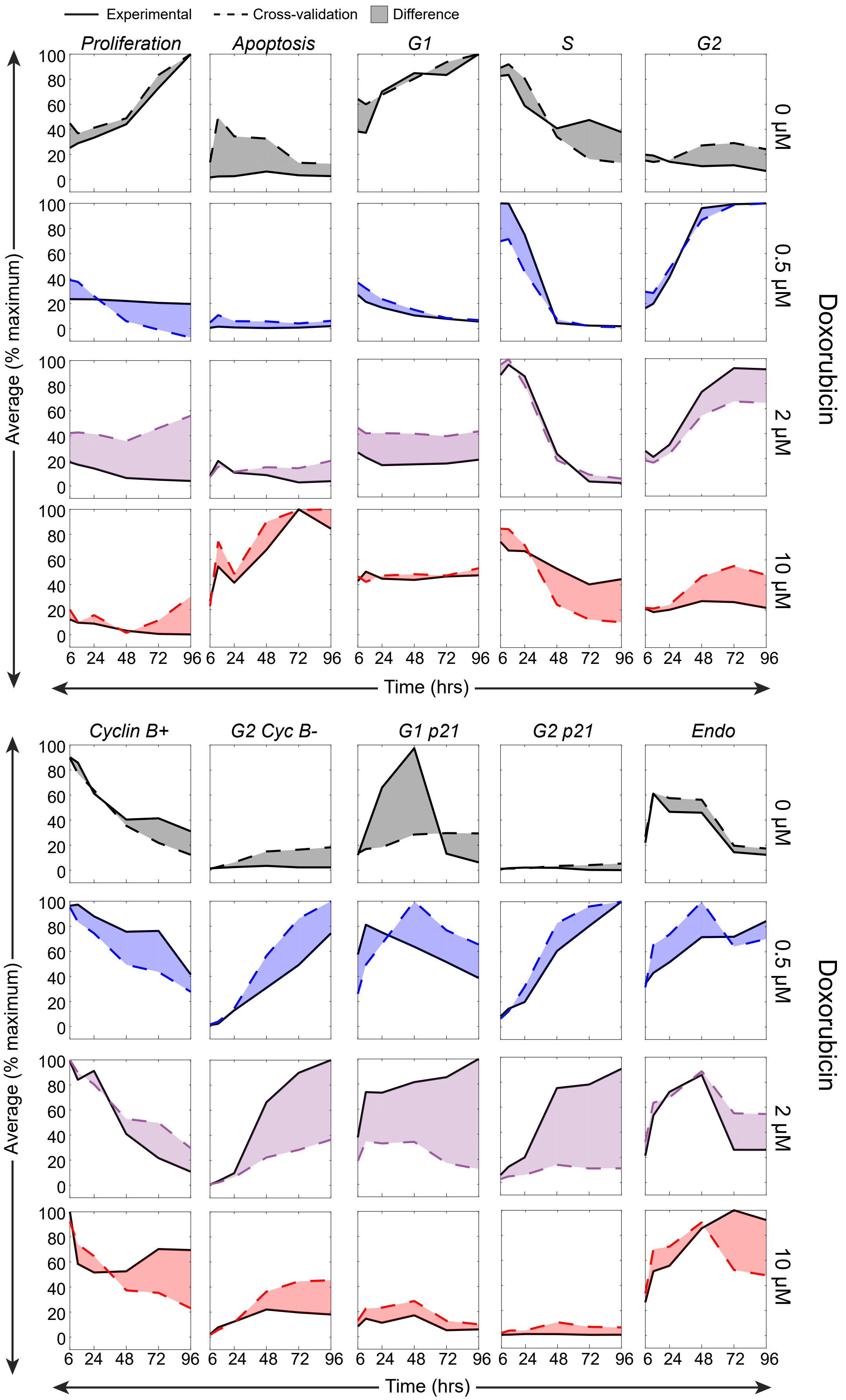

**Figure S4: The 2 μM doxorubicin dose cross-validation values diverge the most from experimental values.**

The mean fluorescence intensity (normalized to the maximum value across time and drug treatments) vs. time is plotted for each of the predicted responses, with the solid lines representing the experimental values, the dotted lines representing the cross-validation predictions, and the shaded area highlighting the difference between the two curves.

### Netterfield et al., Figure S5

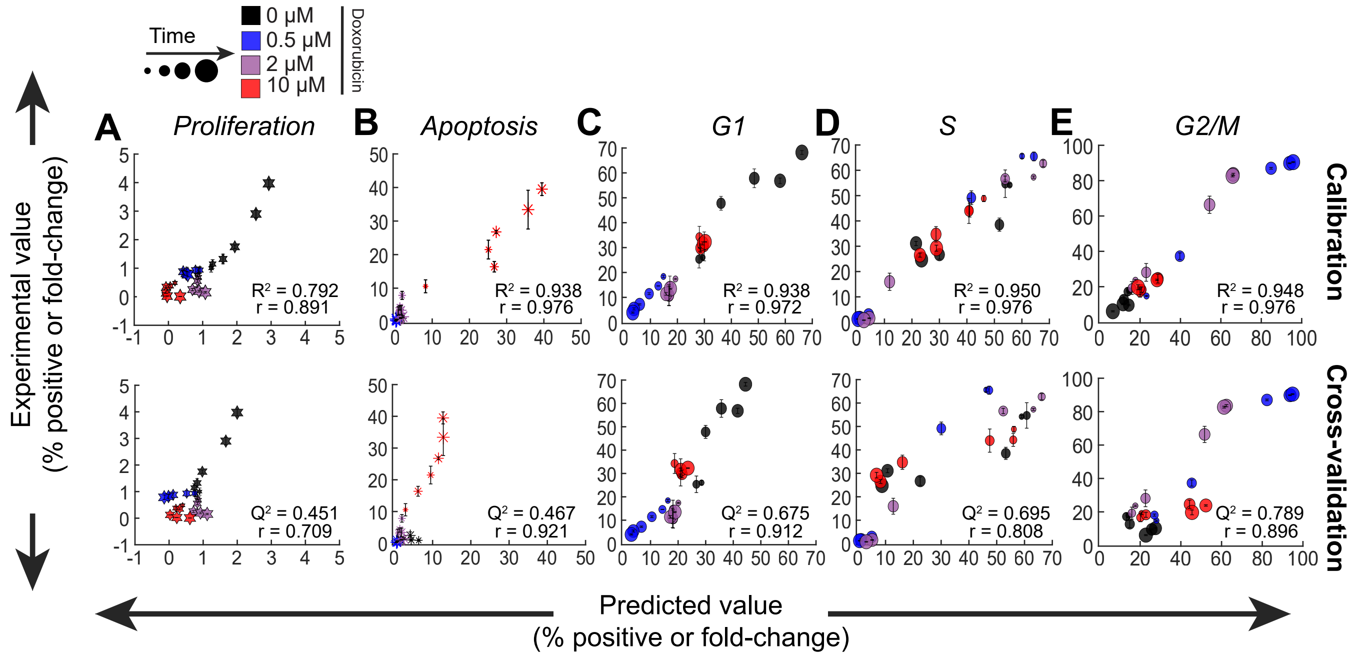

**Figure S5: Experimental vs. predicted values on an individual response basis.**

Scatterplots of experimental vs. predicted values for **A)** Proliferation **B)** Apoptosis **C)** G1 **D)** S and **E)** G2/M responses from the calibration and the cross-validation model. $R^2$, $Q^2$, and Pearson correlation values are also shown.

### Netterfield et al., Figure S6

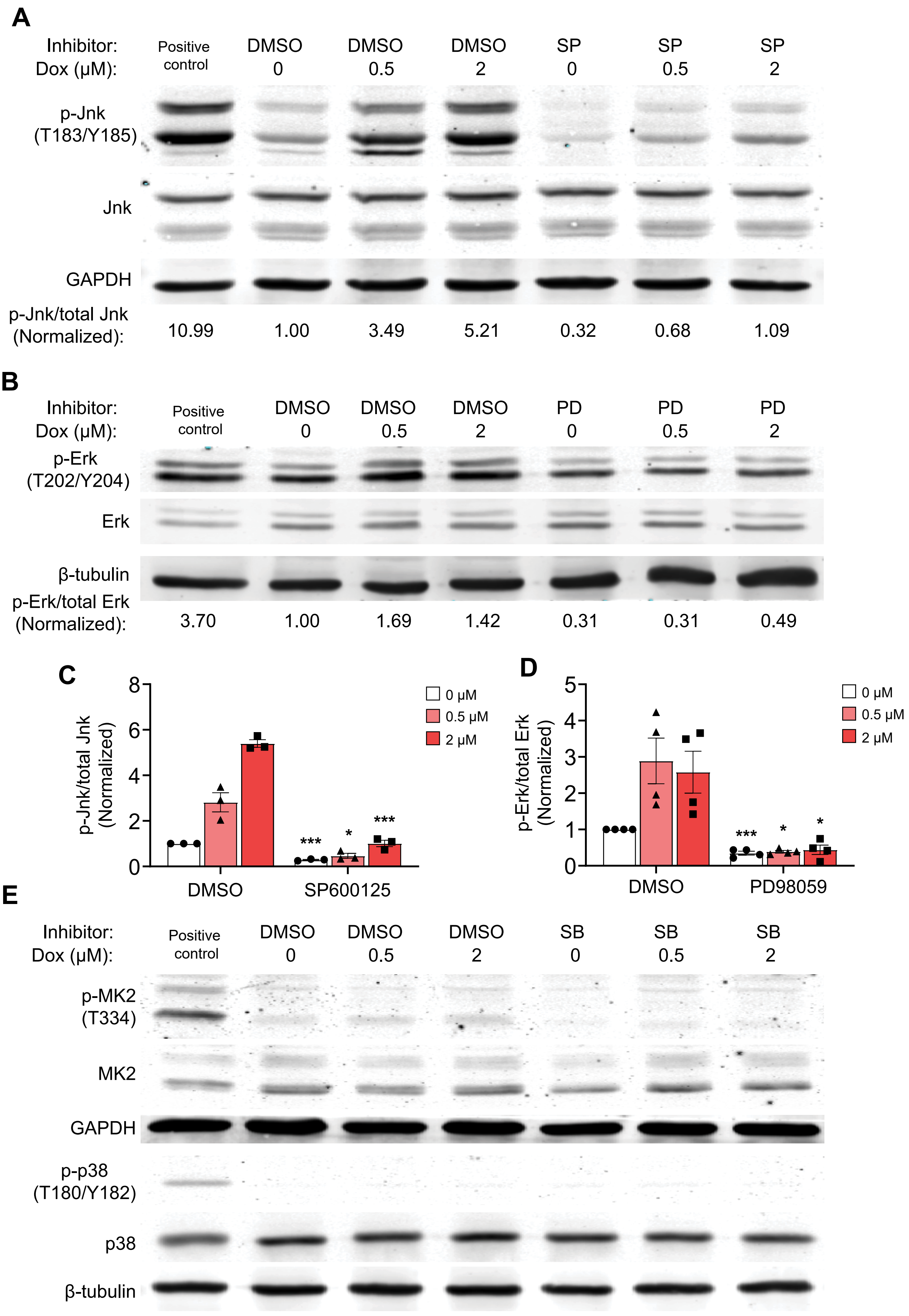

**Figure S6: Western blotting confirms that inhibitors decrease phosphorylation of expected downstream targets.**

**A)** Representative western blot of p-JNK (T183/Y185), JNK, and GAPDH in U2OS cells that were treated with either vehicle (0 µM) or doxorubicin (0.5 µM and 2 µM) for four hours, and cells were either co-treated with vehicle (DMSO), SP600125 (SP, JNKi) or PD98059 (PD, Meki. Samples were run with a positive control of U2OS cells treated with 25 µg/mL anisomycin for 15 minutes.

**B)** Representative western blot of p-Erk (T202/Y204), Erk1/2, and β-tubulin in U2OS samples in the same treatment conditions as subpanel A. Samples were run with a positive control of 100 nM PMA treatment for 30 minutes.

**C)** Quantification of p-JNK to total JNK ratio, normalized to DMSO, 0 µM condition. Bars represent mean ± SEM of three biological replicates.

**D)** Quantification of p-Erk to total Erk1/2 ratio, normalized to DMSO, 0 µM condition. Bars represent mean ± SEM of four biological replicates.

**E)** Representative western blots of p-MK2 (T334), MK2, p-p38 (T180/Y182), p38, GAPDH, and β-tubulin in in U2OS cells that were treated with either vehicle (0 µM) or doxorubicin (0.5 µM and 2 µM) for four hours, and cells were either co-treated with vehicle (DMSO), or SB203580 (SB, p38i). Samples were run with a positive control of U2OS cells treated with 25 µg/mL anisomycin for 15 minutes. Western blots are representative of two biological replicates. ***: $p < 0.001$ *: $p < 0.05$ with a two-tailed t-test with Bonferroni correction tested vs. the respective doxorubicin dose in the vehicle inhibitor control in subpanels C and D.

### Netterfield et al., Figure S7

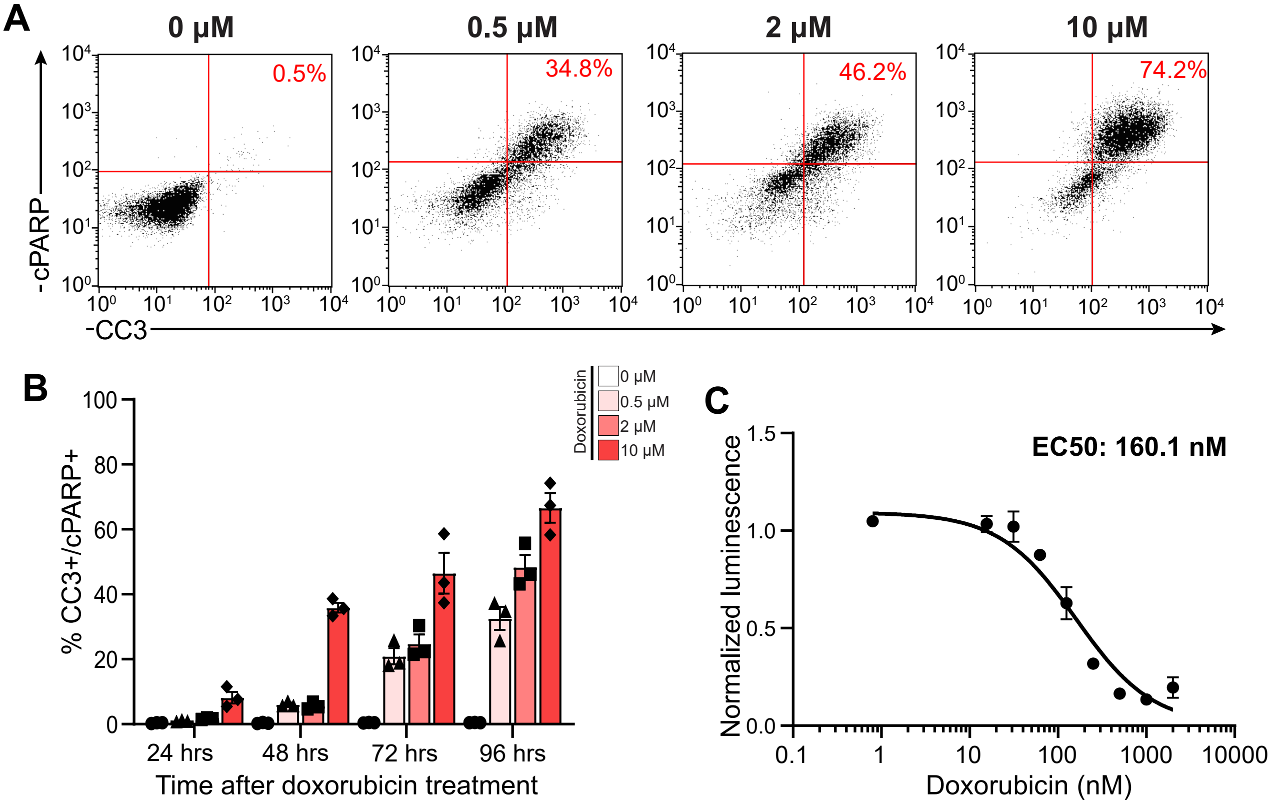

**Figure S7: OVCAR-8 cells undergo dose-dependent apoptosis after doxorubicin treatment**

**A)** Representative flow cytometry plots of cleaved PARP (cPARP) vs. cleaved caspase-3 (CC3) 96 hours after doxorubicin treatment.
**B)** Quantification of CC3/cPARP double positivity over time at varying doses of doxorubicin. Bars represent the mean ± SEM of three biological replicates.
**C)** Viability curve of OVCAR-8 cells using total ATP measurements 96 hours after doxorubicin treatment. All points were normalized to no treatment control, each point represents the mean ± SEM of three biological replicates.

### Netterfield et al., Figure S8

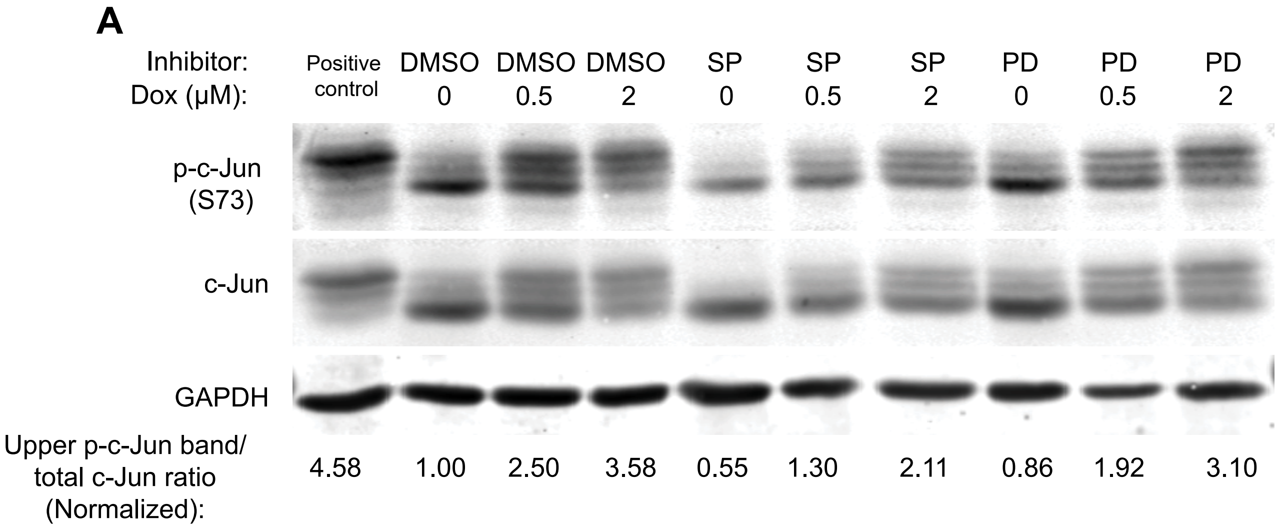


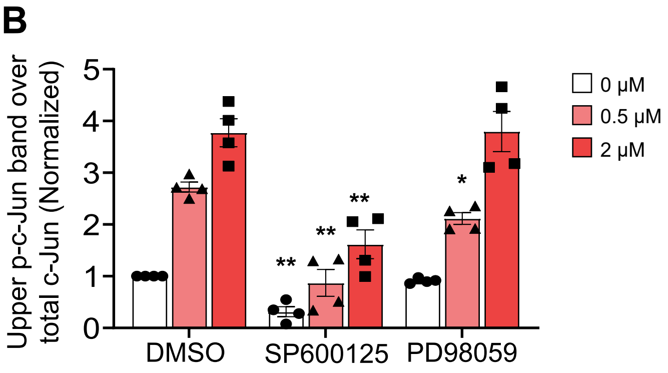

**Figure S8: Immunoblotting reveals a decrease in p-c-Jun(S73) at certain doses of doxorubicin with a JNK or Mek inhibitor co-treatment.**

**A)** Representative western blot probed for p-c-Jun (S73), c-Jun, and GAPDH in U2OS cells that were treated with doxorubicin for four hours, and cells were either co-treated with vehicle (DMSO), or with either 10 µM SP600125 (SP, JNKi) or 10 µM PD98059 (PD, Meki). Cells were lysed 6 hours after doxorubicin treatment for western blotting. Samples were run with a positive control of U2OS cells treated with 25 µg/mL anisomycin for 15 minutes.
**B)** Quantification of the ratio of the upper p-c-Jun(S73) band to total c-Jun. Bars represent the mean ± SEM of four biological replicates, and bars represent SEM. **: $p < 0.01$ *: $p < 0.05$ with a two-tailed t-test with Bonferroni correction tested vs. the respective doxorubicin dose in the vehicle inhibitor control.

### Netterfield et al., Figure S9

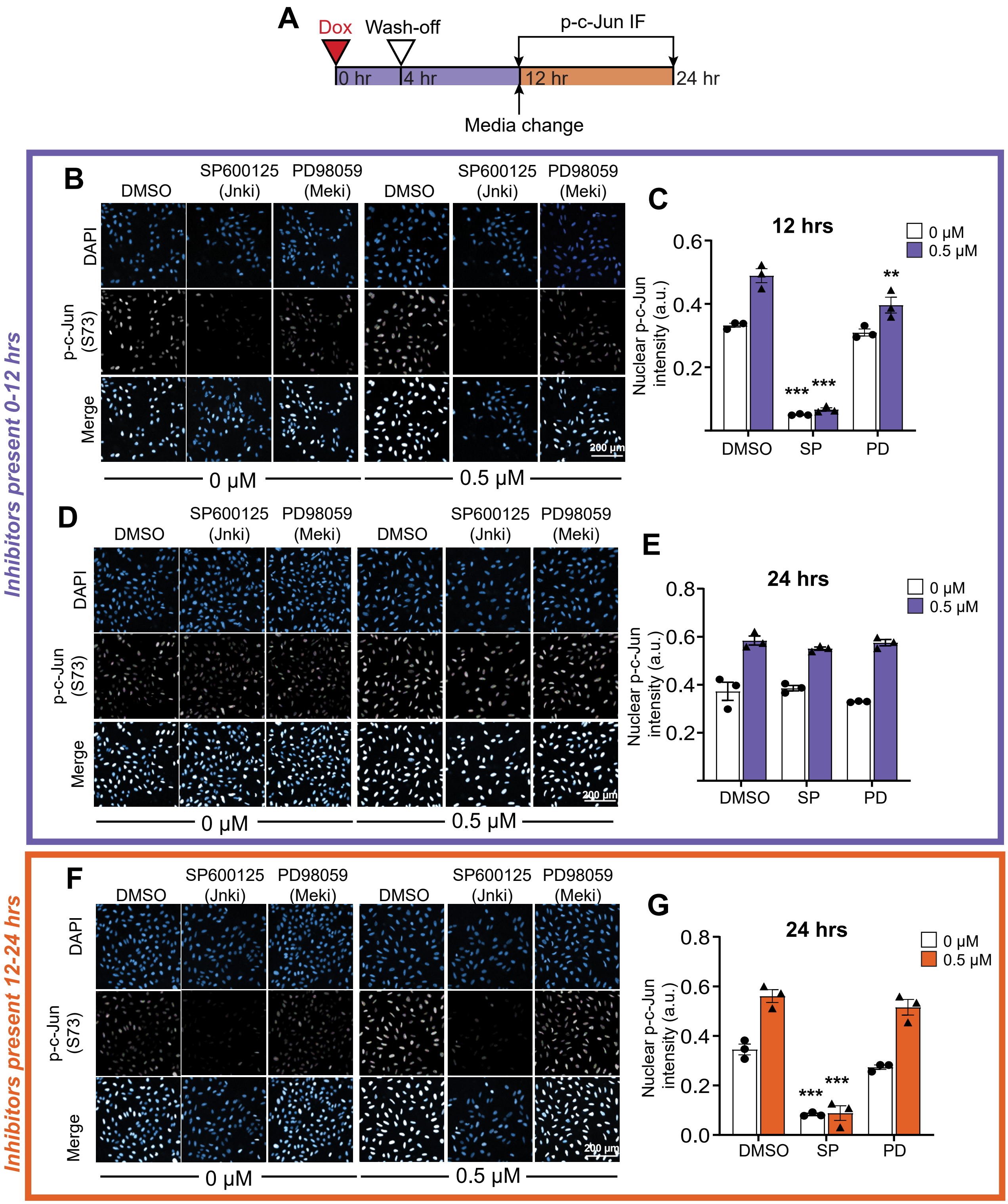

**Figure S9: Time course of p-c-Jun(S73) show inhibitor-dependent decreases in phosphorylation levels in the first 12 hours after doxorubicin treatment in U2OS cells.**

**A)** Schematic of the experiment performed.

**B)** Representative images of cells immuno-stained for p-c-Jun(S73) and DAPI co-treated with inhibitors the first 12 hours of the experiment, and fixed at the 12 hour timepoint.

**C)** Quantification of the nuclear mean p-c-Jun73 intensity 12 hour timepoint.

**D)** Representative images of cells immune-stained for p-c-Jun(S73) and DAPI co-treated with inhibitors the first 12 hours of the experiment, and fixed at the 24 hour timepoint.

**E)** Quantification of the nuclear mean p-c-Jun73 intensity for the 24 hour timepoint after early inhibitor.

**F)** Representative images of cells immune-stained for p-c-Jun(S73) and DAPI co-treated with inhibitors the 12-24 hour window of the experiment, and fixed at the 24 hour timepoint.

**G)** Quantification of the nuclear mean p-c-Jun73 intensity for the 24 hour timepoint after late inhibitor. For subpanels C, E, and G, bars represent the mean ± SEM of three biological replicates. ***: $p < 0.001$ **: $p < 0.01$ with a two-way ANOVA and post-hoc Dunnett's test when compared to the DMSO vehicle control at the same doxorubicin dose.
